# Sorbate Induces Lysine Sorbylation Through Non-Canonical Activities of Class I HDACs to Regulate the Expression of Inflammation Genes

**DOI:** 10.1101/2024.12.11.627973

**Authors:** Yi-Cheng Sin, Breann Abernathy, Douglas G. Mashek, Yue Chen

## Abstract

Metabolic and environmental factors may impact gene expression through the production of active metabolites and epigenetic modifications of histones and transcription factors. In this study, we discovered that cellular uptake of sorbate, an FDA-approved and widely used food preservative, can induce lysine sorbylation (Ksor), a new posttranslational modification and epigenetic mark. We identified over 40 Ksor sites on core histones from mammalian cells and tissue upon sorbate uptake and further showed that the dynamics of histone Ksor could be regulated by the non-canonical activities of Class I histone deacetylases (HDAC1-3). We demonstrated that Class I HDACs catalyzed sorbylation upon sorbate uptake and desorbylation in the absence of sorbate both in vitro and in vivo. Sorbate uptake in mice livers led to a significant increase in histone Ksor without affecting overall histone acetylation, which correlated with the decreased expression of genes in inflammation signaling pathways. Accordingly, sorbate treatment in macrophage RAW264.7 cells upon LPS stimulation dose-dependently downregulated the expression of proinflammatory genes and production of nitric oxide. Global proteomic profiling revealed widespread lysine sorbylation substrates in diverse metabolic and signaling pathways and identified RelA (p65), a component of the NF-ĸB complex, and its interacting proteins as bona fide Ksor targets. Sorbate treatment significantly decreased NF-ĸB transcriptional activities in response to LPS stimulation in RAW264.7 cells. Taken together, our study demonstrated a non-canonical mechanism of sorbate uptake in regulating epigenetic histone modifications and inflammatory gene expression.

## Introduction

Environmental chemicals regulate cellular activities and organismal functions through multifaceted biochemical processes (*1*–*4*). Mechanistic understanding of these processes is instrumental in properly evaluating their physiological impact on human health and developing potential therapeutic interventions. Sorbate is a six-carbon polyunsaturated fatty acid widely used as a chemical preservative in processed food, personal cosmetic products, and pharmaceutical formulations to prevent microbial contamination and ensure product safety (*5*–*7*). Earlier studies in animal and cell models suggested that sorbate uptake had very low cytotoxicity and did not have carcinogenetic activity (*5*–*9*). Yet, more recent studies applying mutagenic and chromosome analysis as well as transcriptome and chromatin analysis showed that sorbate treatment could be potentially genotoxic and affect gene expression in cells and tissues (*10*–*15*). Notably, sorbate treatment may affect fatty acid oxidation pathways and corresponding gene expression in mouse liver (*15*). Thus, the accumulating evidence suggests that chronic sorbate uptake led to changes in gene expression, suggesting a non-trivial role of sorbate uptake in mediating epigenetic dynamics. Given the wide usage of this chemical in food preservation, it is pivotal to uncover the molecular mechanism of sorbate-mediated dynamics of transcription regulation and its potential impact on human physiology.

Gene transcription and cell signaling are intimately linked to chemicals in the cellular microenvironment through protein posttranslational modifications (PTMs) (*16*–*18*). Covalent modifications of proteins through enzymatic or non-enzymatic processes may alter protein structure, enzymatic activity, or protein-protein interaction. Histones and transcription factors are critical targets of active environmental chemicals as the change in their activities profoundly impacts the transcriptional landscape in cells and tissues. With about 500 types of protein modifications reported to-date, over twenty types of modifications have been discovered on histones, and their characterization reveals differential epigenetic effects and new links to cellular metabolism (*19, 20*).

Histone lysine short-chain acylation is a family of protein posttranslational modifications and histone epigenetic marks in mammalian cells (*1*). Starting from lysine propionylation and butyrylation to more recently reported lysine lactylation, these chemical modifications demonstrated that metabolic-linked fatty acids could be activated in situ and served as co-enzymes for acyltransferases that participate in diverse transcription and signaling processes (*21*–*24*). The consequential covalent modification of histones and non-histones has been found to induce protein interactions and regulate enzymatic or transcriptional activities to impact the cellular physiological response to energy and nutrients (*24*–*26*). Inspired by these studies, we showed that the cellular uptake of sorbate, the widely used chemical preservative, induced novel lysine sorbylation (Ksor) in vivo. We demonstrated that Ksor was dynamically regulated by Class I lysine deacetylases (HDAC1-3) that harbor non-canonical functions as either sorbyltransferase or desorbylase depending on the sorbate availability. An increase in Ksor abundance correlated with decreased inflammatory gene expression in mouse liver and cultured macrophage RAW264.7 cells. Global proteomic analysis with immunoprecipitation and LCMS analysis revealed widespread Ksor induced by sorbate treatment and identified RelA (p65), a member of the NF-ĸB complex, as a Ksor target for regulating its transcriptional activities. These studies demonstrated lysine sorbylation as a new histone epigenetic mark and a novel mechanism in sorbate-mediated regulation of inflammation gene expression.

## Results

### Sorbate uptake induces lysine sorbylation, a novel histone epigenetic mark

To determine if sorbate uptake induced endogenous lysine sorbylation, we performed histone extraction and in-gel digestion to sorbate-treated HCT116 human cell line (**figure 1A**). We confidently identified histone peptides with a mass shift of 94.0419 (C6H6O) on lysine residues which matched the expected mass shift of sorbylation on lysine (an example of histone H4 peptide (GLGK(sorbylation)GGAK(acetylation)R was shown) (**figure 1B**). To validate the identified lysine sorbylation, we performed high-resolution LC-MS/MS analysis with HPLC co-elution assay on synthetic peptides with identical sequences and expected modifications as endogenous histone peptides. We observed that the high-resolution MS and MS/MS spectra of the endogenous peptides matched those of the synthetic peptides with very close retention times. Upon mixing synthetic peptides with endogenous peptides, they coeluted perfectly. These data confidently validated the identification of lysine sorbylation in vivo. To further confirm that exogenous sorbate drove lysine sorbylation, we treated the cells with either regular sorbic acid or heavy isotope-labeled ^13^C_2_-sorbic acid (**figure 1C**). The treatment of ^13^C_2_-sorbate led to the identification of ^13^C_2_-lysine sorbylation without the identification of unlabeled Ksor peptides. Comparing precursor ion MS and fragment ion MS/MS spectra of both light and heavy Ksor peptide showed expected mass shifts on precursor ions as well as fragment ions bearing the modification. These data suggested that exogenous sorbate uptake induced histone lysine sorbylation.

**Fig. 1.**
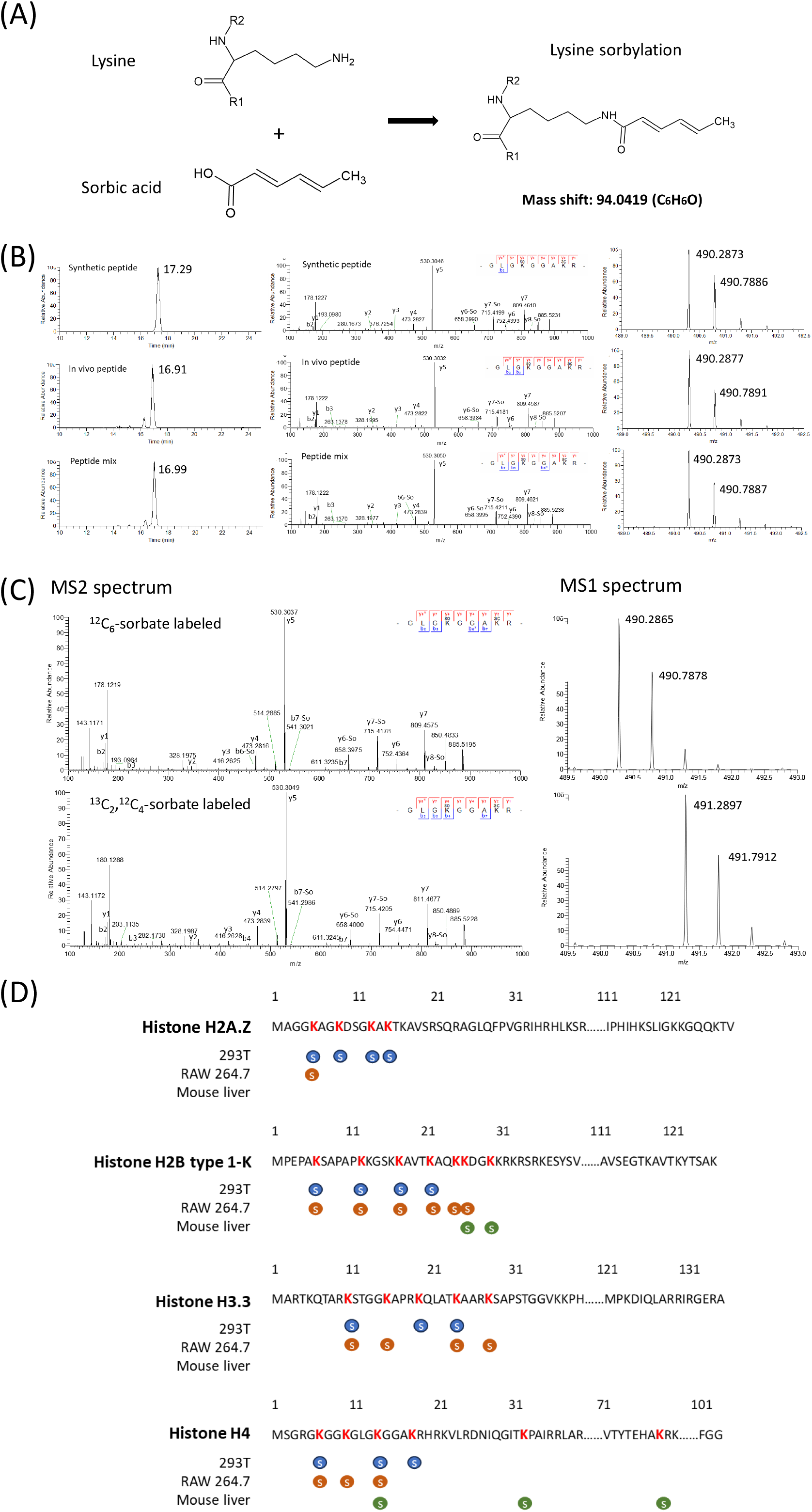
Identification and validation of dietary sorbic acid-induced lysine sorbylation. (A) illustrations for the chemical structures of sorbic acid and the formation of protein lysine sorbylation (Ksor). (B) Extracted ion HPLC chromatograms and MS spectra for the validation of lysine sorbylation with HPLC co-elution analysis. Histone H4 K12 sorbylation peptide “GLGK(So)GGAK(Ac)R” was used as an example (So: lysine sorbylation, Ac: lysine acetylation). “-So” in MS/MS spectra indicates the neutral loss signal of lysine sorbylation. Histones were extracted for HCT116 cells treated with 10 mM of sorbate for 24 hours and subject to SDS-PAGE and in-gel digestion by trypsin. Diluted synthetic peptide, endogenous histone peptides from in-gel digestion and their equal mixture were analyzed with the same HPLC gradient to compare chromatography retention time (left column), MS2 fragmentation (middle column), and peptide MS1 m/z = 490.2804-490.2952 (right column). (C) Validation of lysine sorbylation with isotopic labeling in HCT116 cells. HCT116 cells were treated with 2 mM ^13^C_2_-sorbate for 24 hours and histones were extracted for in-gel tryptic digestion. Peptides were analyzed by LCMS and compared with Ksor peptides from sorbate-treated HCT116 cells. (D) Illustration of representative Ksor sites identified on core histones from the LCMS analysis of histones extracted from sorbate-treated 293T cells (blue), Raw264.7 cells (orange) and mouse liver tissue (green).

To comprehensively identify histone lysine sorbylation, we extracted histones from the sorbate-treated human and mouse cell lines as well as liver tissue from mice that were fasted overnight and then fed diets containing 0.1% and 0.5% (% weight) of sorbate for 12 weeks and performed in-gel digestion for LCMS analysis (**figure 1D**). This level of inclusion in the diet followed the range of 0-25 mg/kg body weight for Acceptable Daily Intake (ADI) for sorbate based on the guidelines from the World Health Organization (WHO) and dose conversion between animals and human (see Materials and Methods) (*27, 28*). We identified a total of 48, 36 and 18 Ksor sites on core histones from 293T cells, RAW264.7 cells and mouse livers, respectively (**figure S1-3 and Data S1**). Interestingly, these histone Ksor sites were located close to the N terminal of the core histones, which overlapped with well-characterized histone acetylation and methylation sites, suggesting a potential role of lysine sorbylation in regulating gene expression through epigenetic mechanisms (**figure 1D**).

To study the dynamic of the lysine sorbylation induced by dietary sorbic acid, we generated a pan anti-sorbylation antibody. We validated the specificity of the Ksor antibody with dot blots against the unmodified peptides, sorbylated peptides, and peptides bearing previously reported acylations with homologous structures, including lysine acetylation and crotonylation (**figure 2A**). Our data suggested that the pan-antibody had high sensitivity and specificity for lysine sorbylation. Using this antibody, we evaluated the induction of histone sorbylation upon sorbate treatment without affecting lysine acetylation in various types of common cell lines, including HepG2, HEK293T, and RAW264.7 (**figure 2B**). Indeed, histone sorbylation was readily induced across different cell lines, suggesting that sorbate uptake could induce lysine sorbylation in cell culture. To determine the temporal dynamics of sorbylation at a whole-cell level, we treated the RAW264.7 cells with 2 mM or 5 mM sorbate at various times. We observed the induction of lysine sorbylation with increasing treatment dosages and treatment times (**figure S4**). To determine if sorbate uptake in mice could induce lysine sorbylation in tissues, we orally gavaged mice with sorbate equivalent to what would be consumed daily with the 0.5% sorbate diet, and harvested mouse livers after various times following the treatment for histone extraction and Western blotting. We observed that the histone sorbylation significantly increased in 2 hours after consuming a diet with sorbate without apparently affecting lysine acetylation compared to control diet (**figure 2C**). This result suggested that the lysine sorbylation could be induced acutely upon sorbate uptake in cell lines and mouse tissues at physiological doses.

**Fig. 2.**
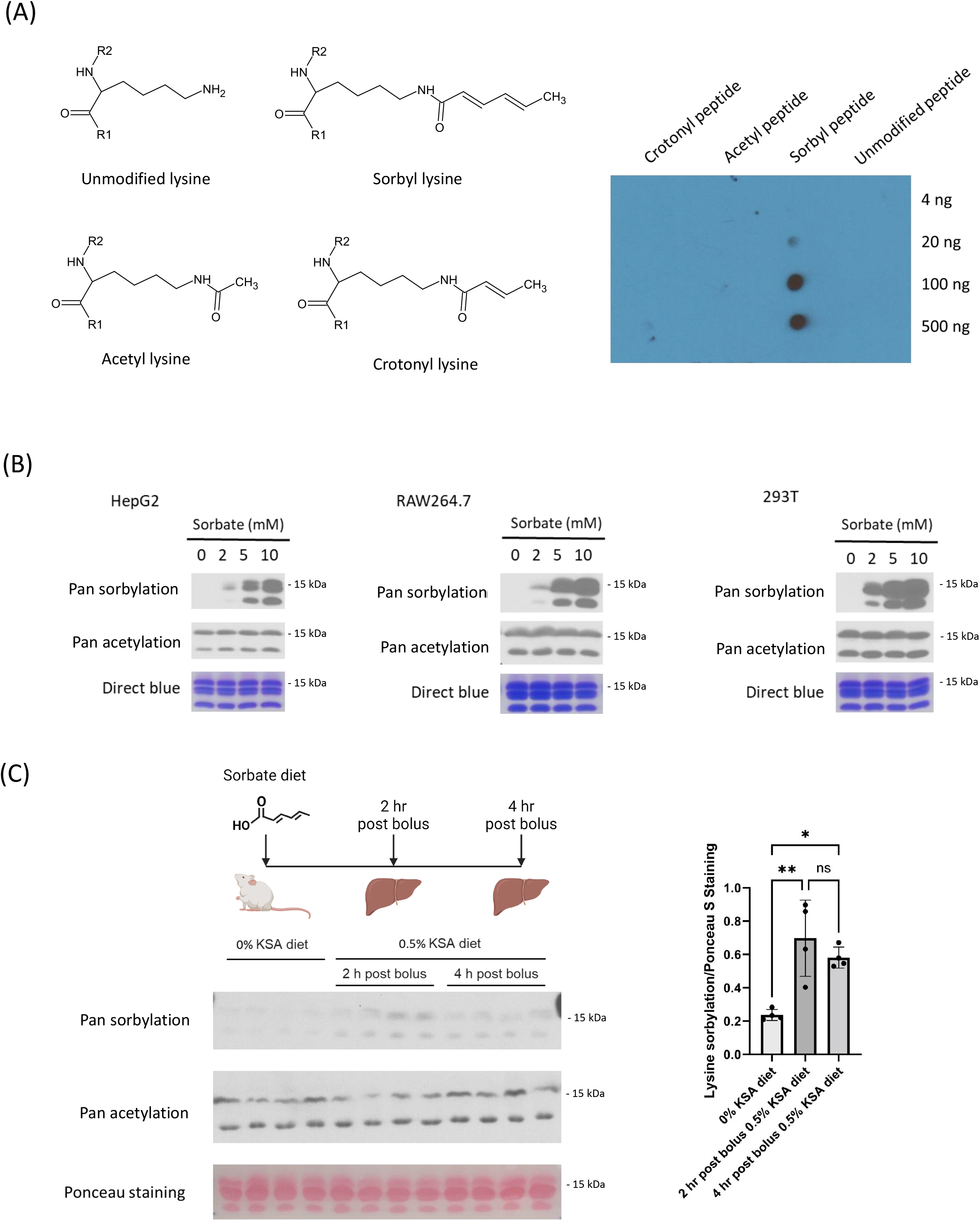
Dynamics of histone sorbylation in cell models and mouse liver tissues. (A) Dot blot assay for the validation of pan anti-Ksor antibody with unmodified peptides or peptides modified with lysine sorbylation, acetylation or crotonylation. (B) Western blot analysis of histones extracted from HepG2 (left), Raw264.7 (middle) and 293T cells (right) treated with various concentrations of sorbate for 24 hours. (C) Western blot analysis of histones extracted from liver tissues of mice fed with a diet containing 0.5% potassium sorbate via oral gavage. Liver tissues were extracted from control mice right away after oral gavage and from mice fed with a sorbate-containing diet 2 hrs or 4 hrs post-feeding. Western blotting was quantified with Image J software and the bar graph quantification was performed with a one-way ANOVA test with Tukey’s multiple comparisons test. Error bars represent standard deviation. * p<0.05 and ** p<0.01.

### Class I histone deacetylase regulates Ksor dynamics

To determine the regulators of Ksor dynamics, we hypothesized that certain members of histone lysine deacetylases (HDACs) known to have promiscuous enzymatic activity could function as lysine desorbylases to remove lysine sorbylation as previously reported for other lysine short chain acylations (*29*–*31*). To this end, we performed a chemical screening with various HDAC inhibitors that have been well-characterized to target different families of HDACs. We expected that the inhibiting HDAC enzymes in cells that functioned as the major lysine desorbylase could retain the level of histone sorbylation after the sorbate removal. To this end, 293T cells were pulsed with sorbate overnight to induce histone sorbylation and then chased with media without sorbate and containing various types of HDAC inhibitors or vehicles. Finally, we harvested cells for histone extraction and western blotting after 5 hours of inhibitor treatments (**figure 3B**). Our data showed that without inhibitor treatments, the abundance of lysine sorbylation diminished quickly in 5 hours (**figure 3B, lanes 2 vs 3**). Treatments with different inhibitors for Class 1, 2, and 4 HDACs retained Ksor levels, but the Class 3 HDAC inhibitor was not as efficient (**figure 3B, lanes 4, 5 vs 6**). Inhibition of class 4 HDAC, which only includes HDAC11, did not retain Ksor (**figure 3B lane 9**). Class 2 HDACs could be classified into Class 2a (HDAC4, HDAC5, HDAC7 and HDAC9) and Class 2b (HDAC6 and HDAC10). Treatment with a Class 2a inhibitor failed to retain histone sorbylation, while the inhibitor towards Class 1 and 2b was highly effective (**figure 3B lanes 7 and 8**). Finally, treatment with MS275 alone, a highly selective inhibitor towards HDAC1, HDAC2, and HDAC3, was sufficient to retain histone Ksor level (**figure 3B lane 10**). We repeated the inhibitor screening in RAW264.7 cells to confirm these findings and observed similar effects (**figure S5**). These data suggested that Class 1 HDACs, including HDAC1, HDAC2, and HDAC3 were potential major histone desorbylases in vivo.

**Fig. 3.**
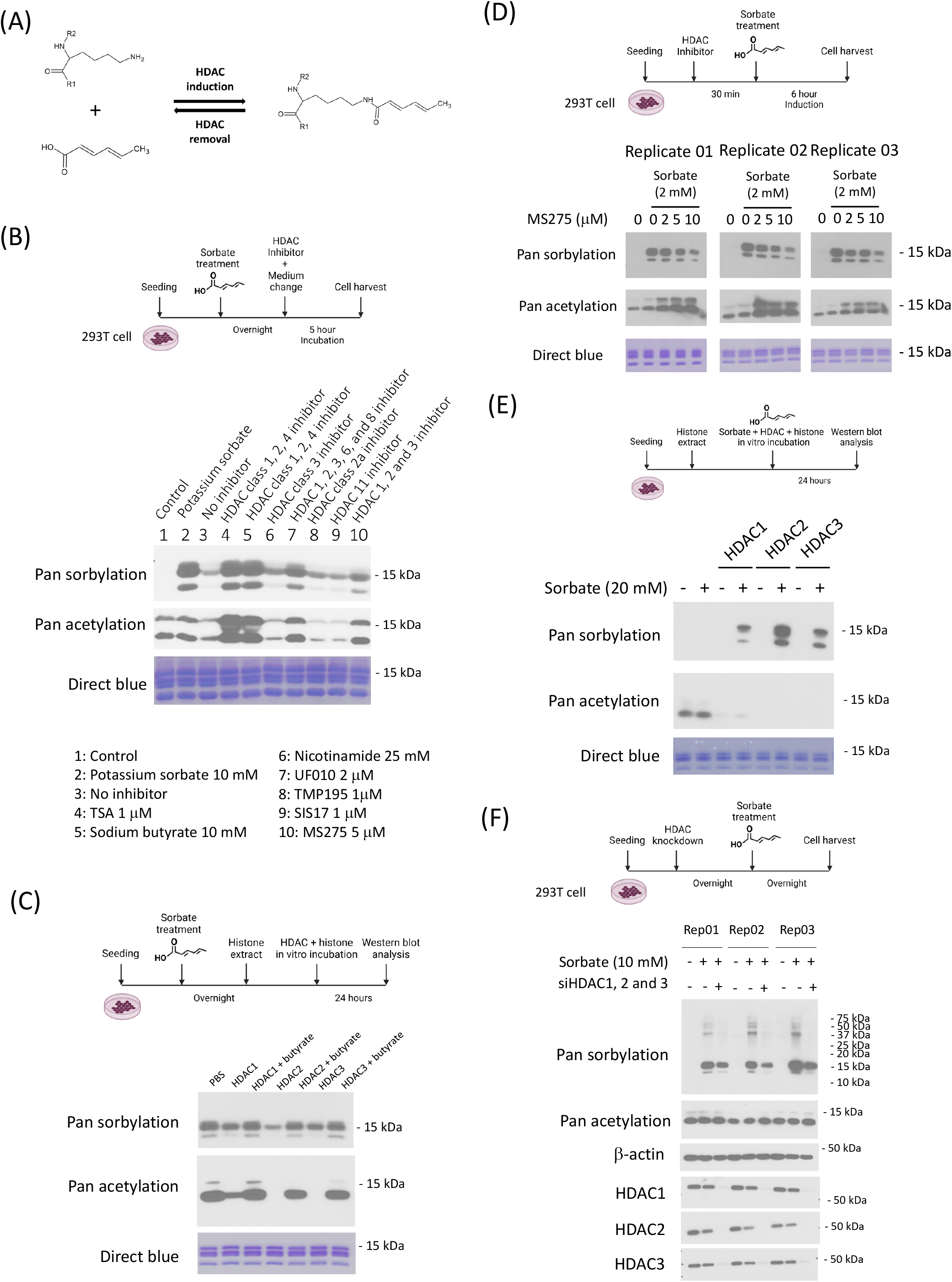
Regulation of lysine sorbylation by Class I HDACs. (A) Schematic representation of HDAC-mediated lysine sorbylation and desorbylation. (B) Chemical inhibitor screening with extracted histones to identify a major class of HDACs that remove histone lysine sorbylation in cells. 293T cells were treated with 10 mM sorbate overnight. Sorbate-containing media were replaced with fresh media with or without various chemical inhibitors for 5 hours targeting different classes of HDACs including Trichostatin A (1 μM, lane 4), Sodium butyrate (10 mM, lane 5), Nicotinamide (25 mM, lane 6), UF010 (2 μM, lane 7), TMP195 (1 μM, lane 8), SIS17 (1 μM, lane 9) and MS275 (5 μM, lane 10) (C) Enzymatic assay and western blot analysis demonstrating the in vitro desorbylase activities of Class I HDACs (HDAC1, HDAC2, HDAC3) with recombinant enzymes and extracted histones from 293T cells treated with or without 10 mM sorbate and 40 mM sodium butyrate for 24 hours. (D) Western blotting analysis shows that HDAC inhibitor MS275 treatment dose-dependently inhibited lysine sorbylation in the presence of sorbate. 293T cells were treated with various concentrations of MS275 first for 30 min followed by the addition of potassium sorbate (2 mM) for the co-treatment of 6 hours before cell harvesting, histone extraction, and western blotting. (E) Enzymatic assay and western blotting analysis demonstrating the in vitro sorbyltransferase activities of Class I HDACs (HDAC1, HDAC2, HDAC3) with recombinant enzymes and extracted histones from regular 293T cells with or without sorbate. (F) SiRNA knockdown and western blot analysis demonstrating the in vivo function of Class I HDACs (HDAC1, HDAC2, and HDAC3) in mediating global lysine sorbylation. SiRNAs targeting HDAC1, HDAC2, or HDAC3 were pooled and transfected to 293T cells for overnight incubation. Media were then replaced with fresh media with or without 10 mM sorbate treatment for overnight. Cells were lysed with SDS sample buffer for western blotting analysis of the whole cell lysate.

To confirm the histone desorbylase activity of these candidate enzymes, we performed in vitro enzymatic assays. Histones extracted from sorbate-treated 293T cells were incubated with equal concentrations of recombinant HDAC1, HDAC2, and HDAC3 enzymes (**figure 3C**). The result showed that all three enzymes exhibited histone desorbylase activities in vitro. The addition of butyrate, a Class I HDAC inhibitor, blocked the enzyme activities of HDAC1, HDAC2, and HDAC3 towards both desorbylation and deacetylation (**figure 3C**). Collectively, our data demonstrated that Class I HDACs are major enzymes that regulate histone lysine sorybylation removal and Ksor dynamics.

An interesting observation was made during our studies that inhibitors of HDACs exerted their effects only when sorbate was removed from the media. Surprisingly, in the presence of sorbate, MS275 (specific inhibitors for HDAC1-3) treatment dose-dependently decreased sorbylation level (**figure 3D**). Inspired by the recent study showing HDAC6 is a lactyltransferase that catalyzes direct lactylation with only lactate as a substrate (*32*), we hypothesized that HDAC1-3 might act as sorbyltransferase in the presence of sorbate. To test this hypothesis, we first performed an in vitro analysis. Histones extracted from cells without sorbate treatment were subject to incubation with recombinant HDAC1, HDAC2, and HDAC3/NCOR2 complex together with or without sorbate, followed by western blotting analysis with pan-anti-Kac and anti-Ksor antibodies (**figure 3E**). These data showed that the enzymes effectively removed acetylation in the absence of sorbate. However, in the presence of sorbate, all three enzymes efficiently installed sorbylation after nearly complete deacetylation. Inhibiting deacetylase activities with TSA treatment also abolished the sorbyltransferase activities of HDAC1-3 (**figure S6**). We further confirmed these findings with TSA treatment in 293T cells and again, our data confirmed that inhibition of HDAC activities abolished sorbate-induced lysine sorbylation while upregulated lysine acetylation in vivo (**figure S7**). These data suggested that HDAC1-3 enzymes harbored non-canonical catalytic activities that functioned as sorbyltransferases in the presence of sorbate without affecting their capabilities as deacetylases and such sorbyltransferase activities were dependent on their deacetylase activities. To determine if HDAC1-3 could function as sorbyltransferases in vivo, we performed siRNA-mediated knockdown of HDAC1-3. Our data showed that the combined knockdown of all three HDACs significantly reduced sorbate-induced lysine sorbylation, suggesting that HDAC1-3 were essential to mediate a significant fraction of lysine sorbylation in vivo (**figure 3F**). These in vivo and in vitro data demonstrated that HDAC1-3 enzymes efficiently catalyzed reversible sorbyltransfer reactions in the presence of sorbate and desorbylation reactions in the absence of sorbate.

### Sorbate uptake regulates proinflammatory gene transcription

To determine how the sorbate uptake may affect endogenous gene expression, we collected mouse liver tissues that showed after 4 hours post bolus treatment of sorbate diet as done in figure 2 and performed RNA-seq analysis. With EdgeR analysis, considering FDR p-value correction and 2X fold change as significant difference gene, we identified 295 genes that were significantly increased in expression and 471 genes showing reduced expression upon sorbate administration (**figure 4A-B, figure S8, Data S2**). Notably, the expressions of numerous inflammation-related genes were down-regulated (**figure 4A**). We performed annotation enrichment analysis of those differentially expressed genes. Indeed, the inflammatory response pathways and related biological processes were significantly enriched among genes with significantly downregulated expression in mice given sorbate (**figure 4C**). This result suggested that sorbate uptake significantly reduced the proinflammatory gene transcriptome in mouse liver.

**Fig. 4.**
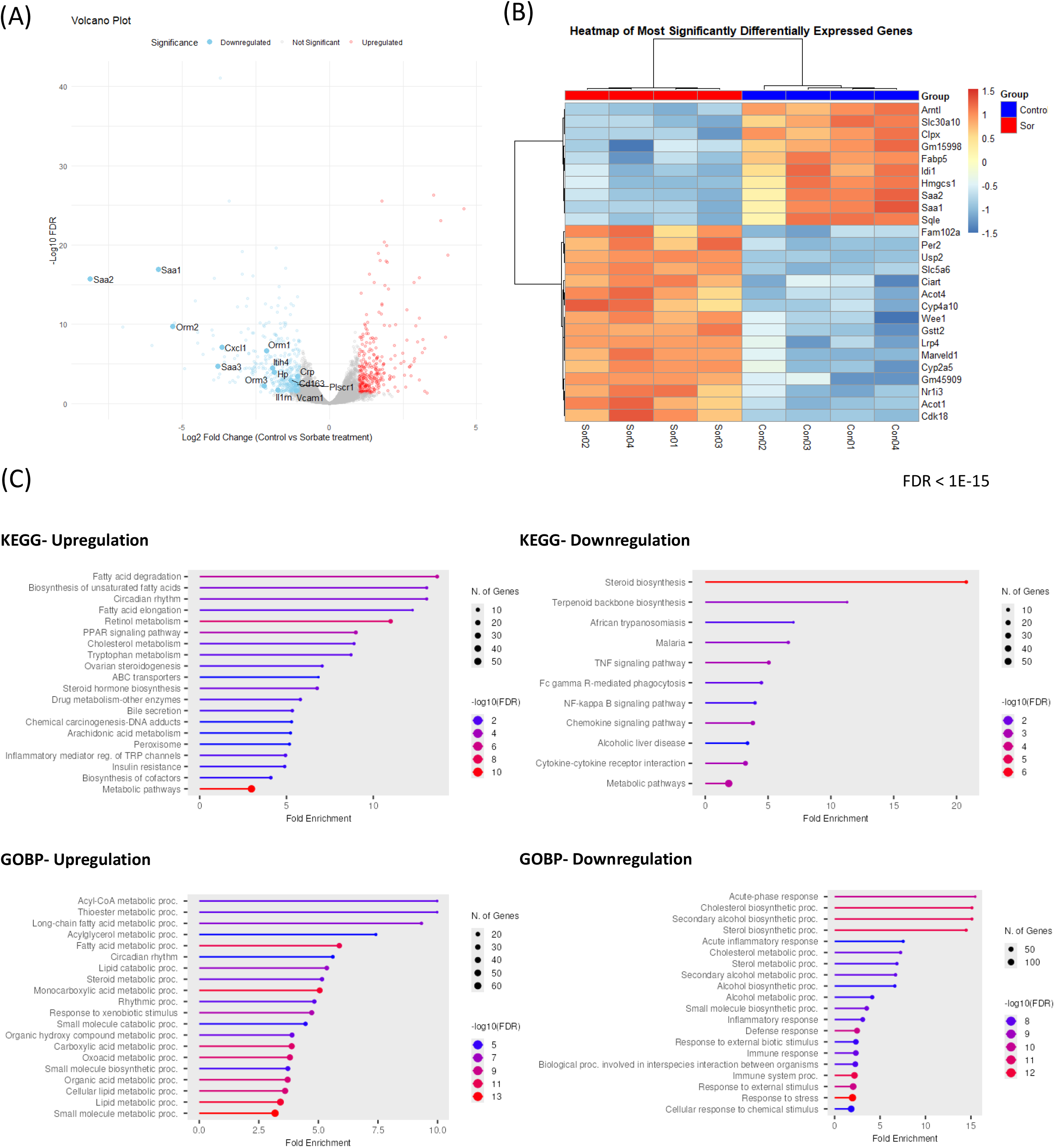
Sorbate-induced transcriptomic dynamics in mouse liver. Liver tissues were extracted from control mice or mice 4 hours post-feeding with 0.5% potassium sorbate in diet via oral gavage. RNAs were extracted for RNA-seq analysis. (A) Volcano plot analysis of gene expression of control and sorbate-treated mouse livers with edgeR (absolute fold change > 2, and FDR < 0.05) representing genes with significant up-regulation (red dots), downregulation (blue dots) and no significant changes (grey dots). Selected genes in acute inflammatory response pathways (GO:0002526) were labeled. Genes with very high y-axis values (-Log10 FDR > 45) were not included in the plot for better visualization. (B) Heatmap and hierarchical clustering analysis of the most differentially expressed genes (FDR < 1e-15) between control and sorbate-treated mouse livers. Scale bar represented standardized (z-score) values (C) Gene ontology annotation enrichment analysis for significantly up-regulated (left column) or down-regulated (right column) genes upon sorbate treatment with KEGG pathway (top row) and Biological Processes (bottom row) analysis. The bar graph represents significantly enriched annotations of genes with a false discovery rate cutoff of 0.05 and absolute fold change > 2.

To confirm these findings, we applied quantitative PCR (qPCR) analysis and studied the expression of inflammatory-related genes in RAW264.7 mouse macrophage cells upon sorbate uptake. To this end, cells were pretreated by various doses of potassium sorbate overnight followed by LPS stimulation for 3 hours (**figure 5A**). qPCR analysis was performed to examine the expression of the proinflammatory genes, including IL6, IL1-beta, IL1-alpha, iNOS and Cox2. Our data showed that sorbate treatment alone did not induce the expression of proinflammatory genes, whereas LPS treatment strongly induced the expression of proinflammatory gene response, as expected. However, when given with LPS, sorbate dose-dependently attenuated the expression of inflammatory genes (**figure 5A**). We further performed nitric oxide assays to determine the effect of sorbate uptake on nitric oxide production. RAW264.7 cells were pretreated with sorbate at various doses overnight, followed by LPS stimulation (**figure 5B**). These data showed that sorbate treatment also dose-dependently reduced the induction of nitric oxide production following LPS. Together, these data demonstrated that the sorbate uptake downregulated proinflammatory gene expression in tissue and cell models.

**Fig. 5.**
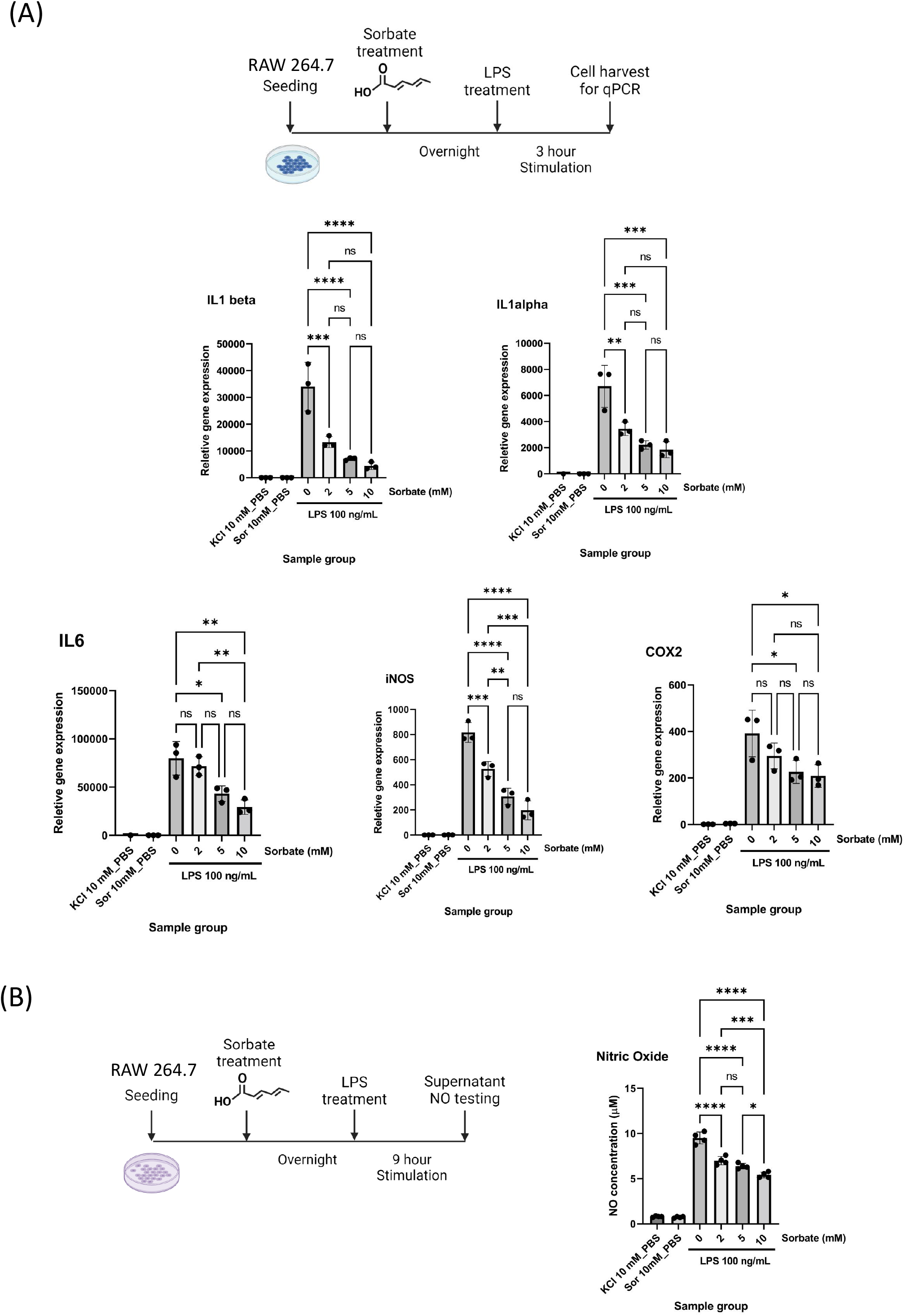
Sorbate treatment downregulated inflammation-related gene expression and NO production in RAW 264.7 cells. (A) Quantitative PCR analysis of relative expressions of IL6, IL-1 alpha, IL-1 beta, iNOS, and COX2 in inflammation response. Cells were treated with various concentrations of potassium sorbate overnight followed by treatment with LPS (100 ng/mL) for 3 hours before cell collection. GAPDH was used as an expression control. (B) Nitric oxide assay to evaluate the regulation of nitric oxide production upon sorbate treatment. Cells were treated with various concentrations of potassium sorbate overnight followed by treatment with LPS (100 ng/mL) for 9 hours before cell supernatant collection and nitric oxide assay. Potassium concentration in each assay was maintained by supplementation with potassium chloride. Quantitative analysis one-way ANOVA test with Tukey’s multiple comparisons test. Error bars represent SD. * p <0.05, ** p <0.01, *** p <0.001 and **** p <0.0001.

### Global proteomic profiling revealed Ksor mediating NF-kB transcriptional activity

To systematically profile global Ksor substrates and identify potential mechanisms for sorbate-dependent anti-inflammatory response in macrophage cells, we performed immunoprecipitation with the pan anti-sorbylation antibody followed by HPLC-MS/MS analysis. Briefly, RAW 264.7 cells were pretreated with 10 mM sorbate overnight and then treated with LPS at 100 ng/mL for 3 hours. Then, the cells were harvested and subjected to lysis and trypsin digestion. The digested peptides were immunoprecipitated with the pan-anti-sorbylation antibody for LCMS analysis (**figure 6A**). We identified over 1600 sites on 908 proteins as Ksor modification substrates (**Data S3**). Bioinformatic analysis showed that Ksor proteins were highly enriched in major cellular pathways including RNA processing, ribosome translation, DNA replication, and cell cycle-related pathways (**figure 6B**). We performed the flanking sequence analysis and observed that lysine sites with serine, proline, and alanine as neighboring residues (+2/-2) were often modified by sorbate, and lysine is the most preferred amino acid beyond two residues surrounding the modification site (**figure 6C**). About 63% of the identified proteins only had one modification site, and nearly 40% were multiply modified with some highly modified proteins bearing more than eight Ksor sites (**figure 6D**). Interaction network analysis identified multiple highly connected Ksor substrate interaction subnetworks related to ribosomal protein translation, pre-mRNA processing, splicing and cell cycle (**figure S9)**. Strikingly, we identified one interaction cluster – RelA (p65) interaction network including p65, Brd4, Stat1, Ncor2, Jun and ATM (**figure 6E**). Given the critical role of p65 and its associated chromatin complex in mediating proinflammatory gene response, these data suggested a potential role of lysine sorbylation in regulating proinflammatory gene expression through modulating p65 transcriptional activities.

**Fig. 6.**
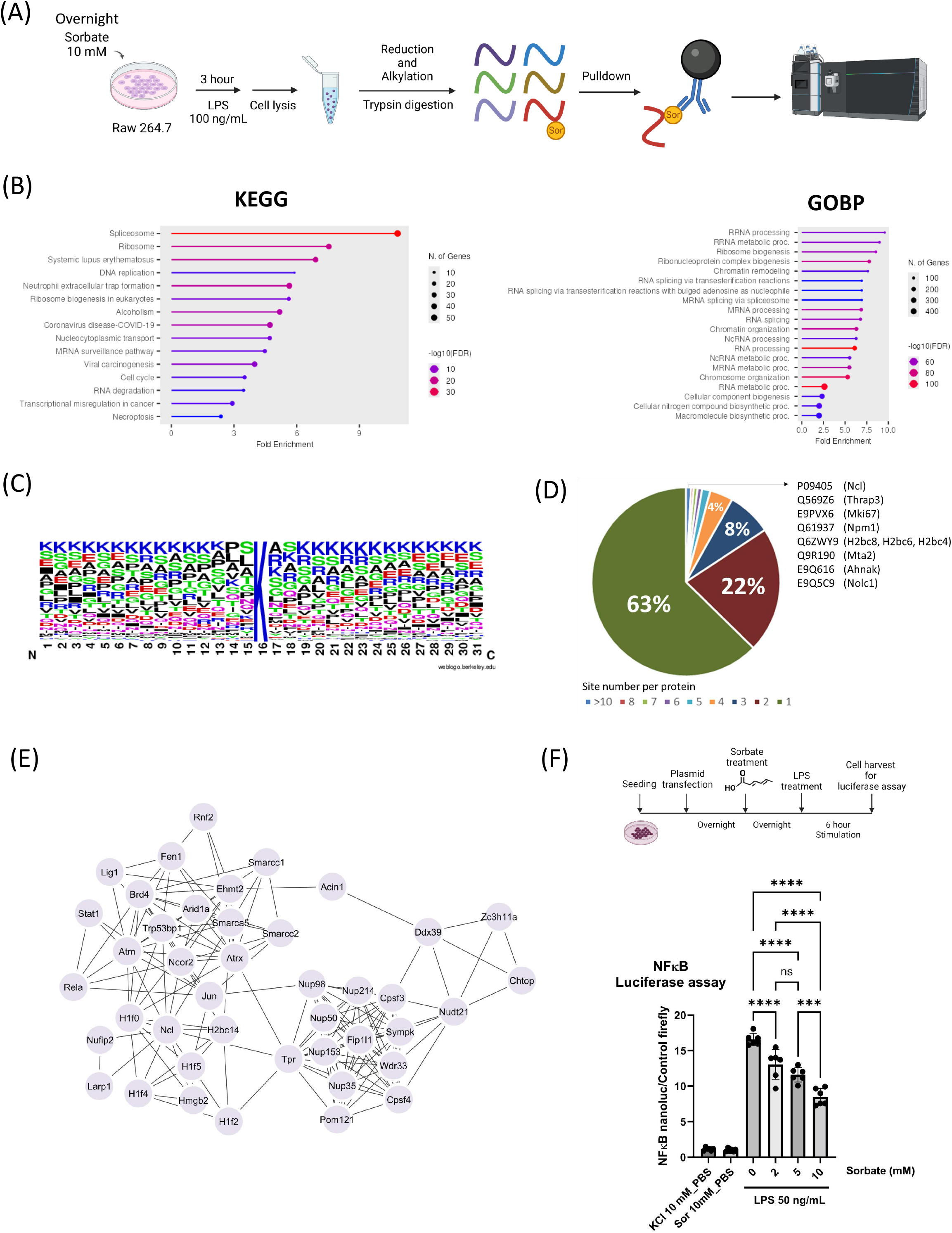
Global profiling of lysine sorbylation substrates and analysis of sorbate-regulation of NF-ĸB activities. (A) Experiment workflow for the immunoprecipitation of lysine sorbylated peptides. Raw264.7 cells were treated with 10 mM sorbate overnight and then stimulated with 100 ng/mL LPS for 3 hours. Cells were lysed for tryptic digestion. Peptides were immunoprecipitated with pan anti-Ksor antibody and identified by Orbitrap LCMS analysis. (B) Annotation enrichment analysis of Ksor substrates with significantly enriched KEGG pathways and Gene Ontology Biological Processes (FDR< 0.05). (C) Flanking sequence analysis of Ksor sites with amino acid abundance within +/-16 positions. (D) Pie chart representing the percentage distribution of site counts in Ksor proteins. (E) Network analysis of Ksor proteins identifying a subnetwork with RelA (p65) interacting proteins. (F) Luciferase assay to monitor the effect of sorbate treatment on Nf-ĸB activity. Nanoluc luciferase plasmid with p65 promoter and firefly control plasmid were transfected to Raw264.7 cells and cells were treated with sorbate for overnight followed by LPS stimulation (50 ng/mL) for 6 hours before cell harvest and luciferase assay. Quantitative analysis one-way ANOVA test with Tukey’s multiple comparisons test. Error bars represent SD. *** p <0.001, **** p< 0.0001 and ns indicates Not Significant (p>0.05).

To determine if p65 transcriptional activity was modulated by sorbate treatment, we performed luciferase reporter assays in Raw264.7 cells. Upon the transfection of luciferase plasmids, cells were treated with sorbate or vehicle control overnight. Then, cells were stimulated with LPS for 6 hours before harvesting for the luciferase assay (**figure 6F**). Our data showed that cellular uptake of sorbate resulted in a significant and dose-dependent decrease in p65 transcriptional activities corresponding to the increasing concentration of sorbate treatments (**figure 6F**). Taken together, our global proteomic profiling identified diverse cellular pathways involved in transcription, translation, and inflammation responses as Ksor substrates and sorbate treatment dose-dependently reduced p65 transcriptional activity in response to the LPS stimulation.

## Discussion

Chemicals in the cellular microenvironment impact physiological activities and organismal functions through diverse mechanisms (*1*–*3*). In this study, we reported that the cellular uptake of sorbate, a widely used food preservative, led to the chemical modification of lysine side chains on proteins in the form of lysine sorbylation. We showed that treatment of mice at a physiological dosage was sufficient to induce sorbylation in the liver. The dynamics of sorbylation were dependent upon sorbate uptake with a relatively quick turnover in several hours (*6*).

Histones were identified and validated as the first lysine sorbylation target. Compared to well-characterized histone acetylation and other short-chain acylations such as propionylation and crotonylation, lysine sorbylation has a six-carbon chain that is more non-polar and has a higher hydrophobicity, indicating a potentially larger impact on the properties and protein interactions of histone proteins in nucleosome as well as with other transcription factors (*21*–*23*). A conjugated double bond offers an electron-rich pi-pi interaction potential that may be targeted by previously reported histone readers for lysine crotonylation and benzoylation (*33*–*36*). Such interaction may result in changes in transcriptional activities and gene expression that alter cellular metabolism and signaling. Continued efforts to discover protein binders that mediate the transcriptional activities of histone sorbylation are important to reveal molecular mechanisms of sorbylation-dependent epigenetic changes in physiology and health.

Through investigating sorbylation regulatory enzymes, we identified Class I HDACs, including HDAC1, HDAC2, and HDAC3, efficiently removed lysine sorbylation and their activities depended on deacetylase activities. Surprisingly, we found that HDAC1, HDAC2, and HDAC3 can act both as sorbyltransferases when sorbate is present and as desorbylases when sorbate is absent, and its sorbyltransferase activities are dependent upon its deacetylase activity. Importantly, we demonstrated that such reversible reactions catalyzed lysine sorbylation independently of the classic CoA activation mechanism in vitro (*10*). Identification of Class I HDACs as both sorbyltransferases and desorbylases is highly significant as it suggests a fast and efficient turnover of lysine acylations from acetylation to sorbylation upon sorbate uptake with an efficient removal of sorbylation in the absence of sorbate. Such reversible catalytic activities of acylation and deacylation are likely not limited to sorbylation and HDAC1-3 enzymes, and the types of acylations with corresponding HDAC enzymes remain to be investigated. Furthermore, the catalytic mechanism of how the enzymes mediate the reversible desorbylation and sorbyltransfer reaction is unclear. Studying these mechanisms will offer new insights into the cellular dynamics of diverse lysine acylation targets in response to changes in fatty acid and energy metabolism. With HDAC1-3 enzymes as catalytic units of several key corepressor complexes, our findings raised intriguing questions on how activities and epigenetic functions of Class I HDACs-related chromatin complexes may be altered upon sorbate uptake and how such changes may impact cellular physiology in human health (*37*).

In our effort to discover the impact of sorbate uptake on gene expression, we performed RNA-seq analysis and discovered that dietary sorbate uptake at a physiological dosage led to a global downregulation of inflammation response pathways in mouse liver. Such findings were confirmed in the mouse macrophage Raw267.4 cell model with qPCR measurements of key proinflammatory genes as well as the production of nitric oxide upon LPS stimulation. While acute inflammation is a key defense mechanism to combat pathogens, chronic inflammation is intimately linked with the development and progression of most metabolic and aging-related diseases (*38*). As such, targeting chronic inflammation is a major focus for mitigating disease development (*39*). Given the novel characterization of sorbate regulating inflammatory pathways, the role of sorbate in disease development warrants further investigation.

Through global proteomic analysis, we identified over 900 proteins as Ksor substrates. We revealed that lysine sorbylation targets diverse cellular pathways in cells with significant enrichment in translation, RNA processing, and cell signaling. Importantly, we identified p65, a key member of the NF-ĸB complex, as a bona fide Ksor target, together with a network of transcription factors and chromatin proteins involved in inflammation response. As a critical mediator of inflammatory response pathways, activities of the NF-ĸB complex are essential for the transcription of proinflammatory genes (*38*). As lysine acetylation is well known to regulate p65 transcriptional activities, it would be intriguing to further study if p65 sorbylation may crosstalk with acetylation and mediate its transcriptional activities (*40*–*42*).

The usage of sorbate has dramatically increased worldwide in the past decade due to the increasing safety demand for processed food, cosmetic, and pharmaceutical products (*43*). In this study, we found that sorbate uptake efficiently induced lysine sorbylation in cells (<5 mM) and mouse tissues (<0.5%) below the level approved by the FDA, suggesting the physiological relevance of lysine sorbylation in regulating organismal metabolism and activities in vivo. Our study identified diverse Ksor substrates and determined their potential role in regulating gene expression and signaling with epigenetic histone modifications and transcription factor targets. As various studies link sorbate consumption with cellular abnormalities and potential health risks, further studies are required to determine the role of lysine sorbylation in normal physiological development as well as diverse pathological situations such as cancers, metabolic diseases, and aging.

## Acknowledgments

We are very grateful for the discussion and suggestions from the members of the Chen lab and Mashek lab, and helpful advice from Drs. David Bernlohr and Timothy Griffin. We greatly appreciate Drs. Yingchun Zhao and Peter Villalta from the mass spectrometry resource center in the Masonic Cancer Center at the University of Minnesota for LCMS setup and access.

## Author contributions

Conceptualization: YS, YC, DM

Methodology: YS, YC, DM

Investigation: YS, BA Visualization: YS

Supervision: YC, DM

Writing—original draft: YC, DM, YS

Writing—review & editing: YC, DM, YS

