## Supporting Information for "Sorbate Induces Lysine Sorbylation Through Non-Canonical Activities of Class I HDACs to Regulate the Expression of Inflammation Genes"

Yi-Cheng Sin *et al.*

**This PDF file includes:**

Materials and Methods
Figs. S1 to S9
Data S1 to S3

### Materials and Methods

#### Materials and Reagents

Lipofectamin 3000 (L3000005), Reverse Transcriptase - SuperScript™ IV VILO™ Master Mix (11756050), Gibco Dulbecco's Modified Eagle Medium (DMEM) (11-965-092), Gibco™ Opti-MEM™ Reduced Serum Medium (31-985-070), TCEP-HCl (PG82080), Laemmli SDS sample buffer (J60015AD), butyrate sodium salt (263191000) and Coomassie Protein Assay Reagent (PI23236) were from ThermoFisher Scientific (Waltham, MA). Sorbic acid (S1626-100G), Luminata Crescenodo HRP substrates (WBLUR0500), polyethyleneimine (PEI), Roche cOmplete protease inhibitor cocktail tablet (04693116001), N-1-naphthylethylenediamine dihydrochloride (NED) (222488), nicotinamide (N3376), sulfanilamide (S9251), fetal bovine serum (F0926), lipopolysaccharide (LPS, L4391), Amicon filter (Ultra-15), MISSION siRNA Universal Negative Control (sic001), siRNAs targeting HDAC1 (SASI_Hs01_00079968) and HDAC2 (SASI_Hs01_00142115) were from MilliporeSigma (St. Louis, MO). siRNA targeting HDAC3 (hs.Ri.HDAC3.13.1) was from Integrated DNA Technologies (Coralville, IA). Recombinant HDAC1 (50051), HDAC2 (50002) and HDAC3/NCOR2 (50003) were from BPS Bioscience (San Diego, CA). Sorbic acid-$^{13}C_2$ (S676891) was from Toronto Research Chemicals (TRC) (Toronto, Ontario, Canada). Trichostatin A (TSA) was from MedchemExpress (Monmouth Junction, NJ). Iodoacetamide (IAA) (02327) was from Chem-Impex (Wood Dale, IL). DharmaFECT 1 Transfection Reagent was from Horizon Discovery (Cambridge, United Kingdom). Monarch Total RNA Miniprep Kit (T1020S) and Luna Universal qPCR Master Mix (M3003) were from New England Biolabs (Ipswich, MA). RNeasy RNA mini kit (74104) was from Qiagen (Venlo, Netherlands). Trypsin (V5280), Nano-Glo Dual Luciferase Reporter (NanoDLR) Assay System with One Glo Reagent and Stop&Glo Reagent (N1610) and luciferase assay plasmids (pNL3.2.NF-κB-RE[NlucP/NF-κB-RE/Hygro] and pGL4.54[luc2/TK]) were from Promega (Madison, WI). 100X penicillin and streptomycin solution (25-512) was from Genesee Scientific (Morrisville, NC). HITRAP protein A resin (17040203) and rProtein A GraviTrap (28-9852-54) were from Cytiva Life Sciences (Marlborough, MA). Empore C18 membrane was from CDS Analytical (Oxford, PA). Mycoplasma PCR Detection Kit (G238) was from Applied Biological Materials (Richmond, British Columbia, Canada). Anti-rabbit IgG HRP-linked antibody (7074S), anti-mouse IgG HRP-linked antibody (7076S), pan anti-acetylated lysine antibody (9441), anti-acetylated-Lysine (Ac-K2-100) MultiMab Rabbit mAb mix (9814), β-Actin (8H10D10) from Cell Signaling (Danvers, MA). Anti-HDAC1 (815102), HDAC2 (680101) and HDAC3 (685201) antibodies were from Biolegend (San Diego, CA). HDAC inhibitors MS275 (Entinostat, HY-12163) and TSA (Trichostatin A, HY-15144) were from MedChemExpress (Monmouth Junction, NJ). SIS17 (S6687), UF010 (S5810) and TMP195 (S8502) were from Selleckchem (Houston, TX). Zinc sulfate (B4382-01, ACS grade) was from Avantor Performance Materials (Radnor, PA).

#### Cell culture

Cells were cultured at 37 °C with 5% $CO_2$ and maintained in Dulbecco's Modified Eagle Medium (DMEM), supplemented with 10% fetal bovine serum, 100 IU penicillin, and 100 μg/ml streptomycin. 293T (CRL-3216), RAW 264.7 (TIB-71), HepG2 (HB-8065) and HCT116 (CCL-247) were from American Type Cell Culture (ATCC) (Manassas, Virginia). Mycoplasma was routinely monitored by Mycoplasma PCR Detection Kit.

### Histone extraction

Cells were scraped down by PBS with 20 mM nicotinamide and 10 mM sodium butyrate. Then, the cell pellets were spun down and lysed by PBS with 0.5% Triton X-100, 1X cOmplete protease inhibitor, 40 mM nicotinamide and 20 mM sodium butyrate. The nuclear was further spun down and extracted by 0.4 N $H_2SO_4$ overnight at 4 °C. Then, proteins were precipitated by 20% (v/v) trichloroacetic acid (TCA) and the precipitated pellet was further washed with ice-cold acetone. To enable SDS gel analysis, the extract was dissolved by non-reducing sample buffer (2% SDS, 10% glycerol, 0.002% bromophenol blue, 100 mM Tris-Cl, pH 6.8).

### Histone in-gel digestion

After the gel was destained, the histone bands were cut, washed thoroughly with water and incubated with 50% ethanol at room temperature with rotation for 1 hour. The gel was further washed with water, cut into pieces and washed with 50% acetonitrile and 100% acetonitrile. Then, the gel pieces were dried down by SpeedVac (Thermofisher Scientific, Waltham, MA). The dried gel pieces were rehydrated with trypsin buffer (1 µg/100 µL of 100 mM ammonium bicarbonate) and incubated at 37 °C overnight. The digestion solution was first transferred, and the peptides in gel pieces were then extracted by 50% acetonitrile and 100% acetonitrile sequentially. Finally, all the peptide solutions were pulled together and dried by SpeedVac and desalted using in-house packed C18 StageTip for LCMS analysis.

### Development of pan-anti lysine sorbylation antibody

Pan-anti lysine sorbylation antibody was developed and purified with the following procedures as previously described (*44–46*). Briefly, lysine-sorbylated keyhole limpet hemocyanin (KLH) was applied to immunize rabbits and serum titer was monitored by ELISA assay (Genemed Synthesis, San Antonio, TX). Pan anti-Ksor antibody was purified from serum using sorbyllysine-conjugated agarose beads and eluted with 0.1M glycine-HCl, pH 2.5, followed by immediate neutralization with 1M Tris-HCl (pH 8.9). The antibody was further dialyzed and concentrated in 1X PBS with an Amicon filter.

### Western blotting

To analyze the whole proteome lysate through western blot, cells were washed with 1X PBS and lysed by Laemmli SDS sample buffer. Then, the lysate was boiled briefly at 99°C with shaking at 800 rpm. Finally, the samples were resolved on SDS-PAGE and proteins were transferred to the PVDF membrane. The membrane was blocked by 5% skim milk or BSA in PBS with 0.1% Tween-20. The membrane was further incubated with primary antibody and HRP-conjugated secondary antibody. Finally, the membrane was developed with Luminata Crescendo Western HRP Substrate for signal detection.

### Dot blot assay

Peptide solutions at various concentrations were dotted onto the nitrocellulose membrane. After the peptide dots were dried, the membrane was blocked with BSA in 1X PBS with 0.1% Tween-20. Then, the membrane was subjected to incubation with primary and HRP-linked secondary antibodies. Signals were developed with Luminata Crescendo HRP substrates.

**Isotopic labeling of lysine sorbylation**

HCT116 cells were treated with 2 mM [$^{13}C_2$]sorbic acid for 24 hours. After the treatment, histones were extracted for SDS-PAGE, stained with Coomassie blue staining and in-gel tryptic digested. The extracted peptides were desalted followed by LCMS analysis.

**HPLC co-elution with synthetic peptides**

Sorbylated peptides were synthesized and purified by GL Biochem Ltd (Shanghai, China) with Fmoc-sorbyllysine (WuXi AppTec, Shanghai, China). The concentration of synthetic peptide solutions was adjusted approximately to match the MS intensities of endogenous sorbylated histone peptides. Endogenous histone peptides, synthetic peptides and their equal mixtures were analyzed through the same HPLC gradient in LCMS analysis with thorough washes in between. Extracted ion chromatograms of mass spectrometry data based on the mass-to-charge ratios of targeted peptides were presented.

**Assay of nitric oxide production in RAW264.7**

RAW264.7 cells were seeded in 24-well plates overnight. Then, the cells were treated with sorbate for 15 hours. After the sorbate treatment, LPS was given to cells for the other 9 hours of incubation. 100 µL of supernatant from the cell was tested by mixing with 50 µL of 0.1% N-1-naphthylethylenediamine dihydrochloride in water and 50 µL of 1% sulfanilamide in 5% phosphoric acid. Sodium nitrite dissolved in water was used as standard curve construction. Finally, the product was detected by absorption spectroscopy at 545 nm.

**HDAC knockdown**

293T cells were seeded in 12-well plates. siRNA (0.05 nmol each) targeting human HDAC1, HDAC2 or HDAC3 was mixed with DharmaFECT 1 transfection reagent in DMEM medium and incubated with cells for 30 hours, while the control cells with sorbate treatment were added with siRNA Universal Negative Control (0.05 nmol) mixed with DharmaFECT 1 transfection reagent in DMEM medium and the control cells without sorbate treatment were added only with DharmaFECT 1 transfection reagent in DMEM medium. After the incubation, the original medium was replaced with fresh DMEM medium and subjected to treatments with sorbate or PBS.

**HDAC in vitro assay**

*Desorbylase Activity Assay* Histones extracted from 293T cells treated with 10 mM sorbic acid for 24 hours were reconstituted by the assay buffer (PBS with 1 mM $MgCl_2$, 10 µM zinc sulfate and 10 mM Tris, pH 7.5-8.5) to a final concentration of 0.5 mg/mL. Then, histones were incubated with an HDAC enzyme at a final concentration of 500 nM with or without 40 mM butyrate for incubation with rotation at 37 °C for 24 hours. Finally, reactions were stopped by boiling in a non-reducing sampling buffer (2% SDS, 10% glycerol, 0.002% bromophenol blue, 100 mM Tris-Cl, pH 6.8) for western blot and Coomassie blue staining analysis.

*Sorbyltransferase Activity Assay* Histones extracted from regular 293T cells were reconstituted in the assay buffer as above to a final concentration of 0.5 mg/mL. Then, histones were incubated with an HDAC enzyme at a final concentration of 500 nM with or without various concentrations of potassium sorbate or TSA (50 µM) for incubation with rotation at 37 °C for 24 hours. Finally, reactions were stopped by boiling in the non-reducing sampling buffer for western blot and Coomassie blue staining analysis.

**Immunoprecipitation of Ksor peptides**

Raw264.7 cells were seeded in 15 cm plates and treated with 10 mM potassium sorbate overnight. Then, the cells were stimulated by LPS 100 ng/mL for 3 hours. The treated cells were scraped down in PBS and lysed in denaturing lysis buffer (9 M Urea with 1X cOmplete protease inhibitor), followed by reduction and alkylation with TCEP and IAA at the final concentration of 10 mM at room temperature in dark. Then, the reaction was quenched by cysteine (20 mM). The cell lysate was diluted to 1.5 M Urea with 100 mM ammonium bicarbonate and trypsin was added at an enzyme-to-substrate ratio of 1:50 (w/w) for proteolytic digestion at 37 °C overnight. The digested peptides were desalted and dried in SpeedVac (Thermofisher Scientific, Waltham, MA). The dried peptides were dissolved in the IP buffer (1X PBS with 0.1% Tween-20) and incubated with protein A beads pre-conjugated with pan anti-sorbylation antibody overnight. The beads were washed with the IP buffer four times and 1X PBS four times. Finally, the peptides were eluted by 0.1% TFA three times. The eluted peptides were desalted using in-house packed C18 StageTip for LCMS analysis.

**Conversion of sorbate ADI between human and mouse**

Acceptable Daily Intake (ADI) of sorbate for man is 25 mg/kg body weight (*27*). Based on the formula below (*28*)

Animal equivalent dose (AED) (mg/kg) = Human dose (mg/kg) / Km ratio

(Km ratio is the correction factor, and the value is 0.081 to convert the human dose to the mouse dose)

From this calculation, we obtained the sorbate ADI for mice as 308.6 mg/kg. If we assume the reference body weight of mouse is 25 g or 0.025 kg (*47*), then we obtained the equivalent dose of sorbate uptake per day as 7.7 mg. If we assume mice eat 4 grams of food per day, the percentage of sorbate in food in mouse diet is approximately 0.2%.

**Treatment of mice with a diet containing potassium sorbate via oral gavage**

Twelve mice (C57BL6/J males ~10 weeks of age) were given either a control (water) bolus (n=4) or 17 mg of potassium sorbate via oral gavage. This dosage represents the approximate average daily intake of potassium sorbate around 0.5% of the diet during the feeding trial. The control mice were sacrificed immediately, and sorbate mice were sacrificed 2 or 4 hours after gavage (n=4/group).

**Mouse liver RNA extraction and sequencing**

RNA was extracted from snap-frozen liver tissue using the RNeasy RNA mini kit. Eukaryotic RNA isolates were quantified using a fluorometric RiboGreen assay. The total RNA integrity was assessed using capillary electrophoresis. Only samples higher than 1 μg with a RIN of 8 or greater proceeded to sequencing. Total RNA samples were converted to Illumina sequencing libraries using Illumina’s Truseq RNA sample preparation kit. One microgram of total RNA was oligo-dT purified using oligo-dT-coated magnetic beads, fragmented, and then reverse transcribed into cDNA. The cDNA was fragmented, blunt-ended, and ligated to indexed (barcoded) adaptors and amplified using 15 cycles of PCR.

The final library size distribution was validated using capillary electrophoresis and quantified using fluorimetry (Pico Green) and via quantitative (q)PCR. Indexed libraries were then normalized, pooled, and size selected to 320 bp using a Caliper XT instrument. Truseq libraries were hybridized to a single read flow cell, and individual fragments were clonally amplified by bridge amplification on the Illumina cBot. Once complete, the flow cell was loaded on the Hi-Seq 2500 and sequenced. Upon completion of read 1, an 8 bp forward and 8 bp reverse (i7 and i5) index read was performed. Base call files for each cycle of sequencing were generated by Illumina Real Time Analysis (RTA) software. Primary analysis and demultiplexing were performed using Illumina bcl2fatstq software version 2.17.1.14.

Data alignment and gene quantification were analyzed using the CHURP pipeline at the University of Minnesota Supercomputing Institute (MSI). 2 x 150bp FASTQ paired-end reads for 12 samples (26.1 million reads average per sample) were trimmed using Trimmomatic (v0.33) enabled with the optional "-q" option; 3bp sliding-window trimming from 3' end requiring minimum Q30. Quality control on raw sequence data for each sample was performed with FastQC. Read mapping was performed via HISAT2 (v2.1.0) (*48*) using the mouse genome (GRCm38) as a reference. Gene quantification was done via Feature Counts for raw read counts. Differentially expressed genes (DEGs) were analyzed by R with the edgeR library using raw read count as input. The volcano plot and heatmap figures were generated with the libraries including ggplot2, ggrepel, pheatmap, dplyr, and annotables. We filtered the generated list based on a minimum absolute fold change > 2, and FDR corrected $p < 0.05$.

**qPCR assay**

RAW264.7 cells were treated with various concentrations of potassium sorbate overnight followed by LPS at 100 ng/mL for 3 hours before harvesting. The total RNA was extracted by Monarch Total RNA Miniprep Kit and subjected to reverse transcription with SuperScript IV VILO Master Mix according to the manufacturer's instructions. Finally, qPCR was performed with Luna Universal qPCR Master Mix and the primers below.

1. Mouse GAPDH primers:

Forward sequence: CATCACTGCCACCCAGAAGACTG

Reverse sequence: ATGCCAGTGAGCTTCCCGTTCAG

2. Mouse IL6 primers:

Forward sequence: TACCACTTCACAAGTCGGAGGC

Reverse sequence: CTGCAAGTGCATCATCGTTGTTC

3. Mouse PTGS2 primers:

Forward: GCGACATACTCAAGCAGGAGCA

Reverse sequence: AGTGGTAACCGCTCAGGTGTTG

4. Mouse iNOS2 primers:

Forward: GAGACAGGGAAGTCTGAAGCAC

Reverse sequence: CCAGCAGTAGTTGCTCCTCTTC

5. Mouse IL1a primers:

Forward sequence: CGAAGACTACAGTTCTGCCATT

Reverse sequence: GACGTTTCAGAGGTTCTCAGAG

6. Mouse IL1b primers:

Forward sequence: GCCACCTTTTGACAGTGATGAG

Reverse sequence: GACAGCCCAGGTCAAAGGTT

**LCMS analysis**

Peptides were analyzed with the Dionex Ultimate 3000 RSLC nano HPLC system with an Orbitrap Fusion Lumos Tribrid Mass Spectrometer (ThermoFisher Scientific, Waltham, MA) similarly as previously described (*49*). Briefly, digested peptides dissolved in HPLC buffer A (0.1% formic acid in water (v/v)) were separated on an in-house packed capillary HPLC column (length 20 cm and 75 µm inner diameter) from CoAnn Technologies (Richland, WA) with Luna C18 beads (5 µm particle size, 100 Å pores) from Phenomenex (Torrance, CA). The peptides were separated with a gradient from 1% to 95% HPLC buffer B (0.1% formic acid in acetonitrile (v/v)) in HPLC buffer A. Full scan of precursor ions (MS1) was performed in a positive mode in Orbitrap and MS/MS analysis was performed in the linear ion trap with a 35% of normalized high-energy collision dissociation (HCD) energy.

**Luciferase assay**

Raw 264.7 cells were seeded in 12-well plates overnight and then transfected with NF-κb Nanoluc plasmid and control Firefly plasmid with Lipofectamine 3000 based on manufacturer's protocol. After overnight transfection, media was replaced with fresh DMEM and cells were treated with various concentrations of potassium sorbate or potassium chloride overnight. Then, cells were treated with LPS for six hours. After the LPS stimulation, media was removed, and cells were briefly rinsed with PBS followed by lysis with passive lysis buffer. The protein lysate was centrifuged at 21,000 ×g for 5 min at 4 °C to remove pellets. ONE Glo Reagent was added to the supernatant of the lysate to measure firefly luminescence and Stop&Glo reagent was added to measure the second luminescence.

**Database searching**

LCMS data was analyzed with the Maxquant database search engine (version 2.2.0.0) (*50*). Uniprot mouse protein database (UP000000589_10090) was searched for the LCMS analysis of Ksor immunoprecipitation from Raw264.7 cells. A small database containing human and mouse histone proteins from the UniProt protein database was searched for the LCMS analysis of extracted histones. Trypsin was specified as the protease and the maximum number of missing cleavages was set as 4. Cysteine carbamidomethylation was set as a fixed modification for Ksor immunoprecipitation sample analysis. Methionine oxidation, protein N-terminal acetylation, lysine acetylation and lysine sorbylation were defined as variable modifications. Sorbylation modification was defined as C(6)H(6)O and neural loss was set as C(6)H(6)O. The maximum number of modifications per peptide was set as 4.

**Bioinformatics**

For protein cluster interaction analysis, Ksor protein interaction network was extracted from the STRING database with default criteria and exported for presentation with Cytoscape (*51*, *52*). Subnetworks of Ksor protein clusters were identified with MCODE analysis (*53*). Annotation enrichment analysis of DEGs for KEGG pathway and Gene Ontology Biological Processes was done using ShinyGo (ver 0.81, FDR < 0.05) (*54*). Flanking sequence analysis was performed with Weblogo (version 3) with +/-16 positions of Ksor sites (*55*). Additional R packages were used for programing including readxl, dplyr and openxlsx. Graphpad prism (GraphPad Software, Boston, MA) was used for bar graph representation with statistical analysis. Figure preparation was assisted by Biorender. Chemical structures were drawn with ChemSketch 2022.2.2 version.

**Fig. S1. Example spectra of Ksor sites identified from histone extract in sorbate-treated 293T cells.** 293T cells were treated with 10 mM potassium sorbate for 24 hours and histones were extracted for in-gel tryptic digestion and LCMS analysis.

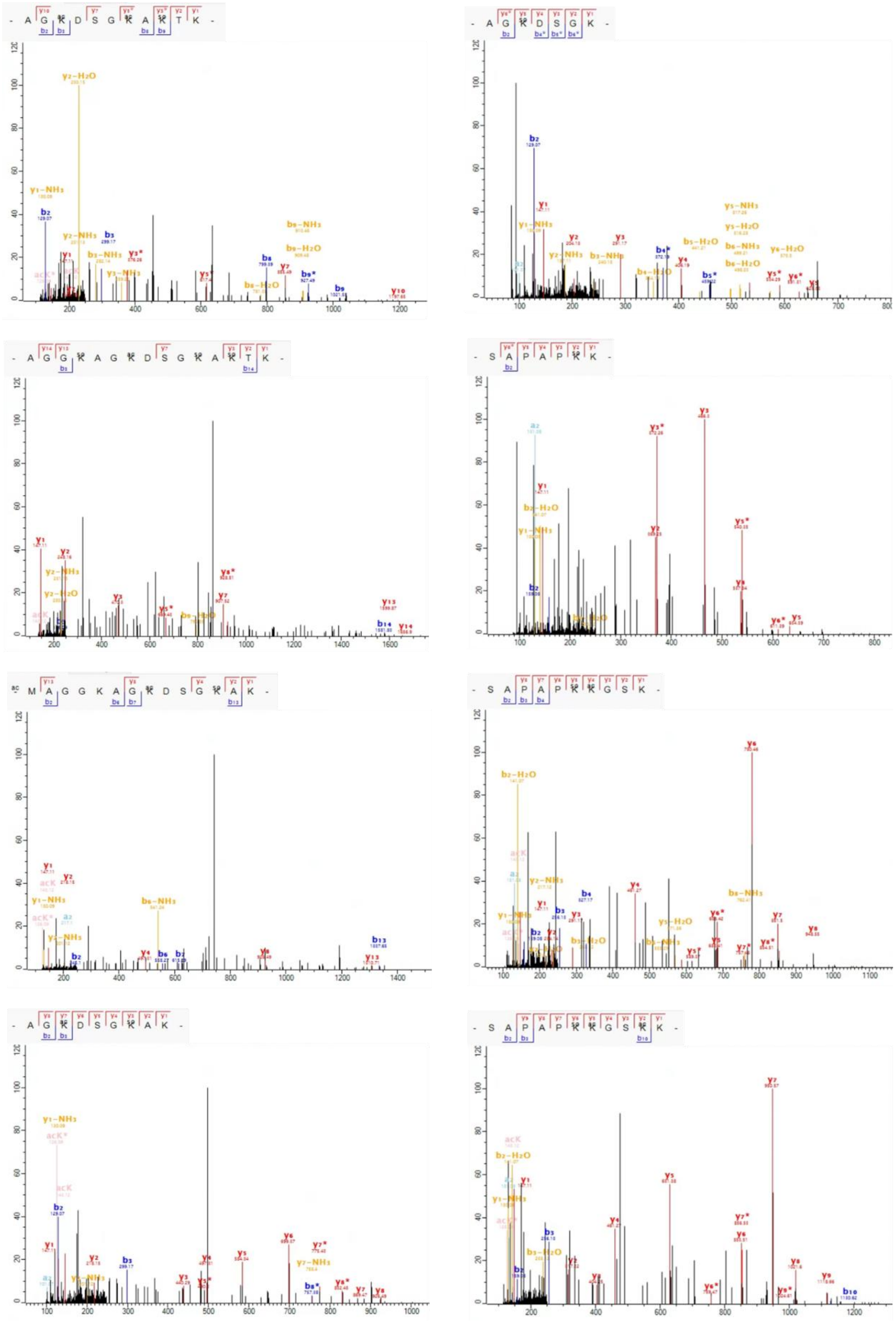

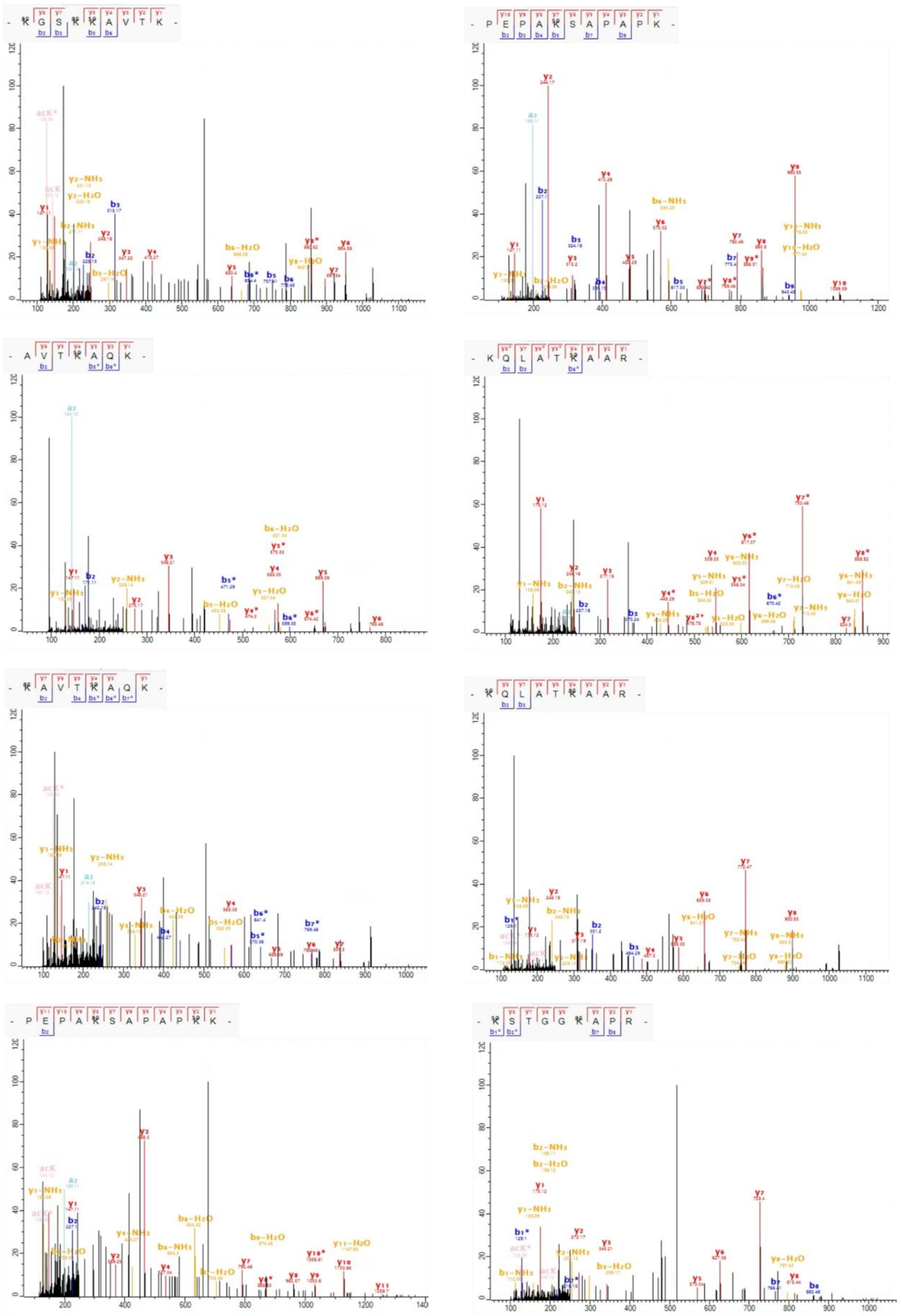

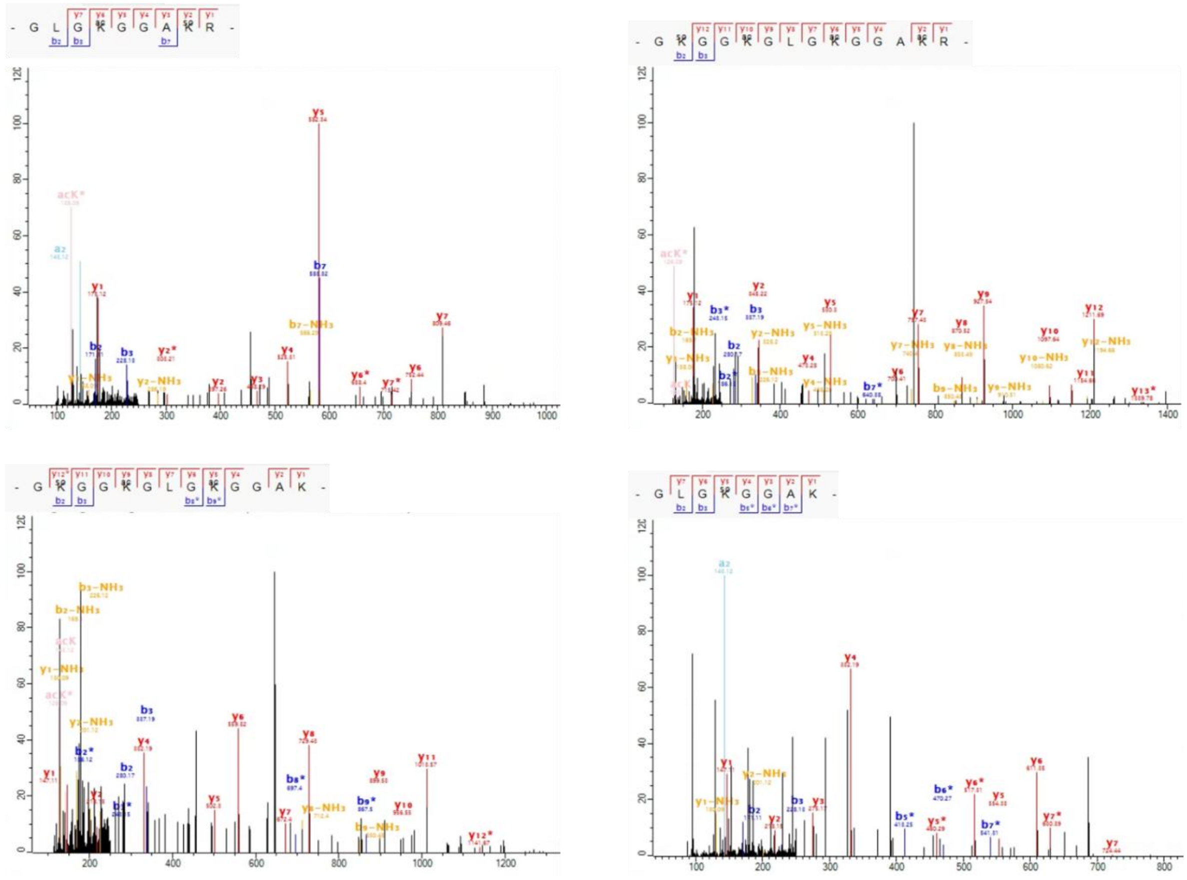
- G L G K G G A K R -
- G K G G K G L G K G G A K R -
- G K G G K G L G K G G A K -
- G L G K G G A K -

**Fig. S2. Example spectra of Ksor sites identified from histone extract in sorbate-treated RAW 264.7 cells.** RAW 264.7 cells were treated with 10 mM potassium sorbate for 24 hours and histones were extracted for in-gel tryptic digestion and LCMS analysis.

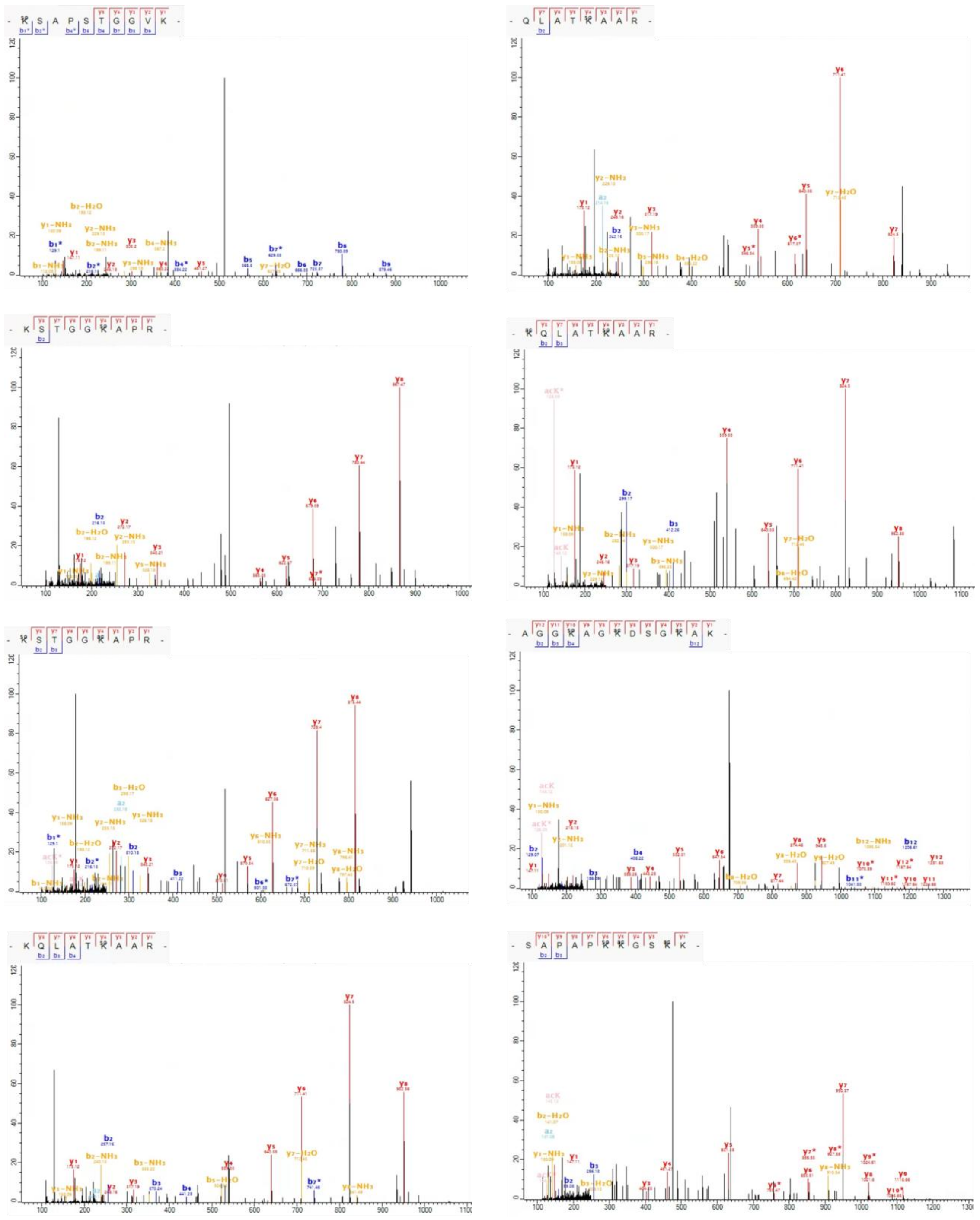

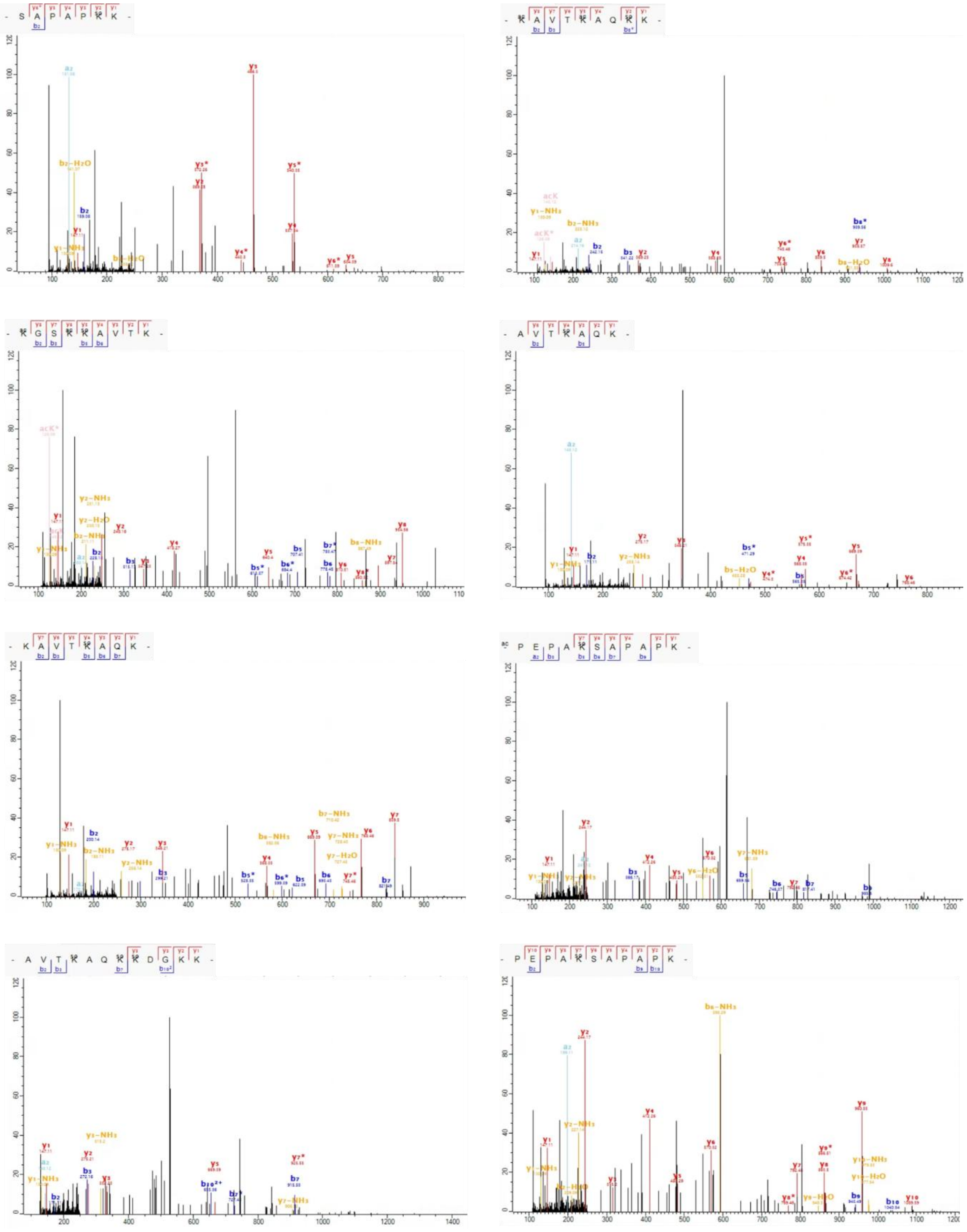

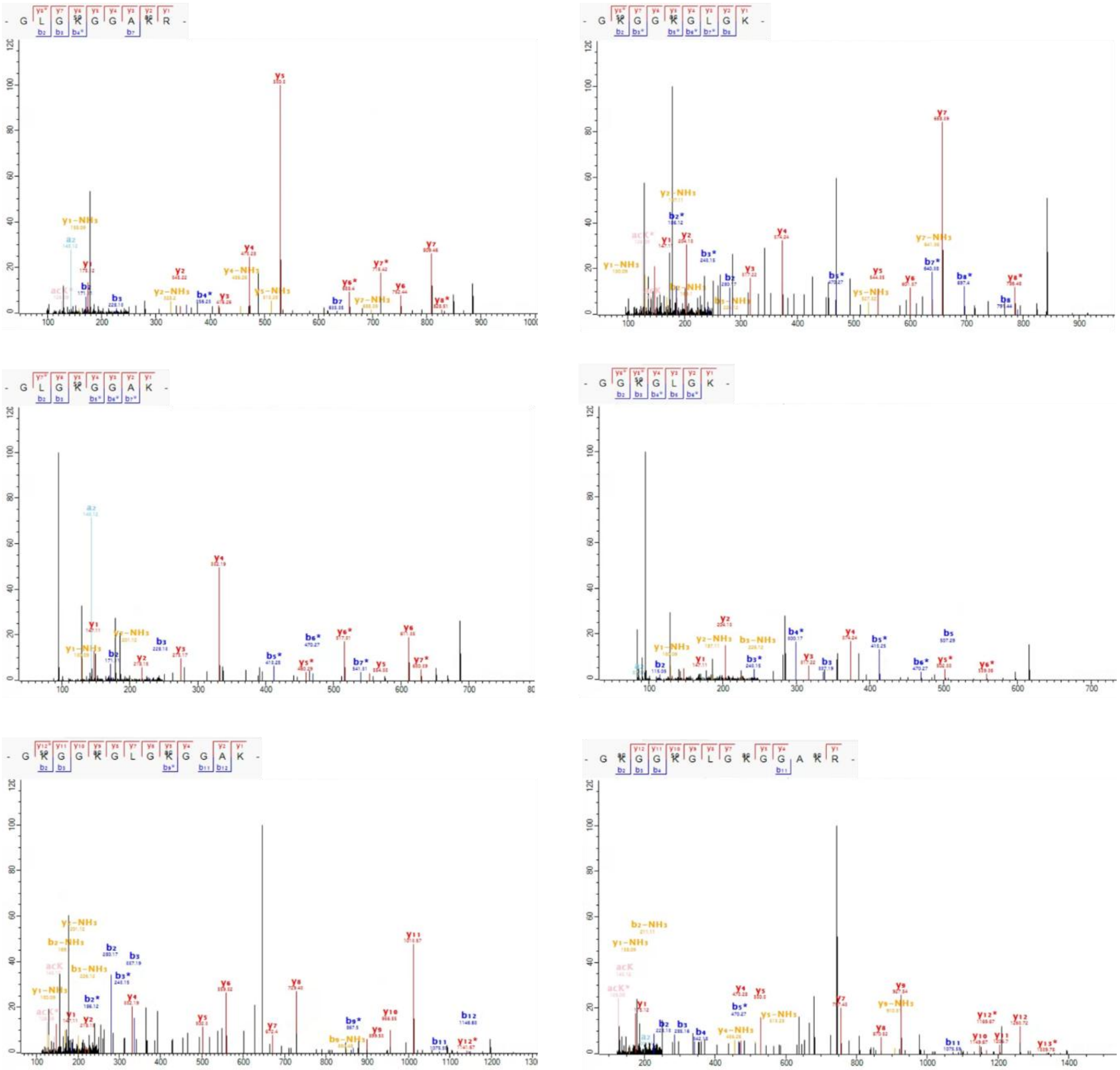

**Fig. S3. Example spectra of Ksor sites identified from histone extract in liver from mice fed with diet containing 0.1% or 0.5% (%weight) sorbate for 12 weeks.** Histone proteins were extracted for in-gel tryptic digestion and LCMS analysis.

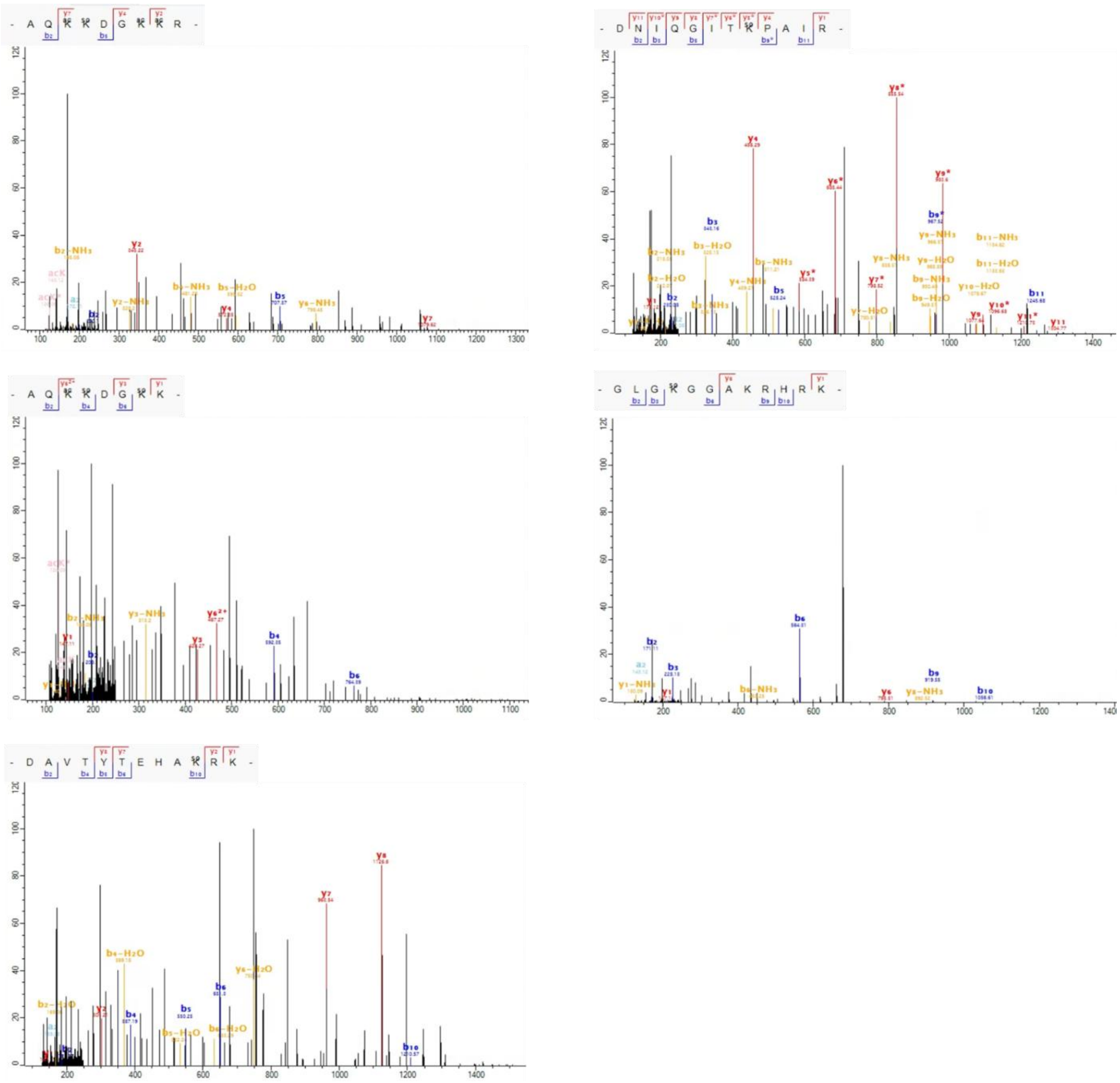

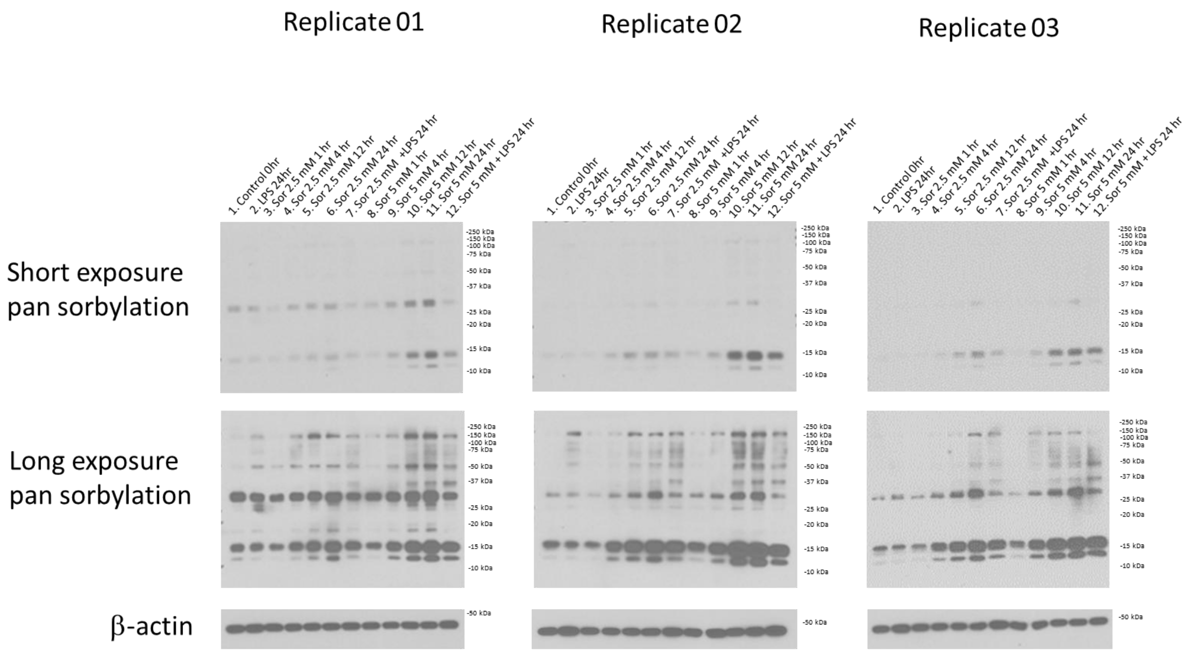


**Fig. S4. Western blotting analysis of Ksor dynamics in the whole cell lysate of RAW264.7 cells.** RAW264.7 cells were treated with or without sorbate or LPS (100 ng/mL) in various dosage and time prior to cell lysis and western blotting with pan anti-Ksor antibody.

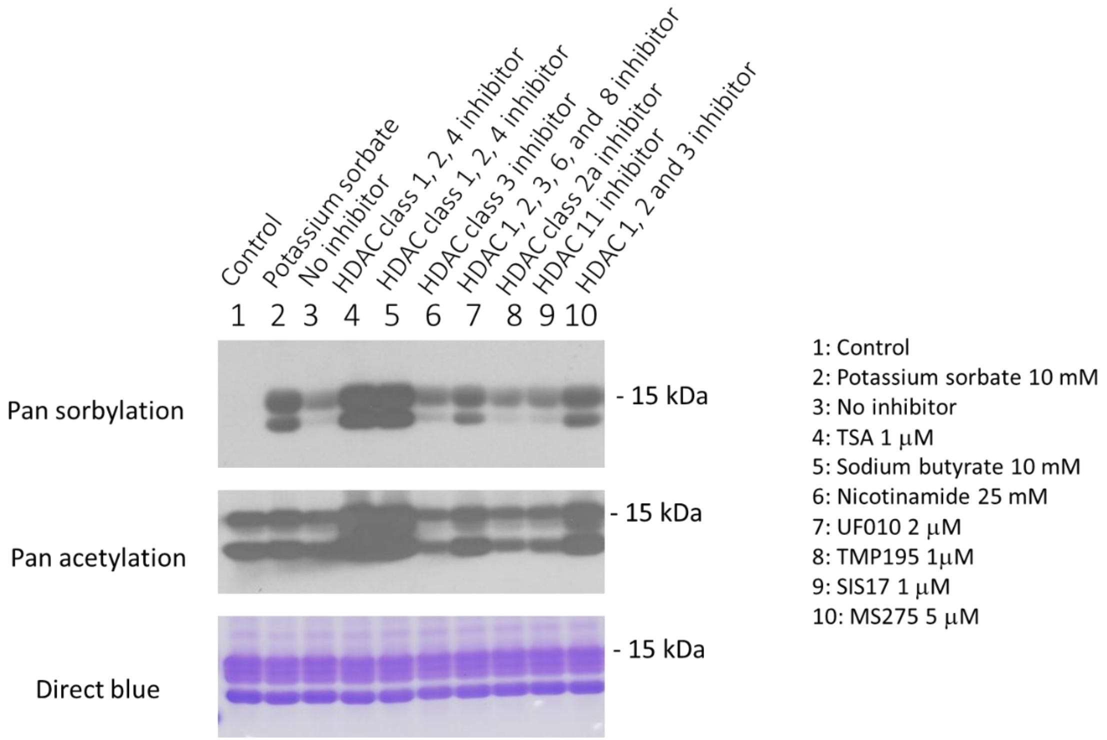


**Fig. S5. Chemical inhibitor screening with extracted histones to identify major class of HDACs that remove histone lysine sorbylation in RAW264.7 cells.** Raw264.7 cells were treated with 10 mM sorbate overnight. Sorbate-containing media were replaced with fresh media with or without various chemical inhibitors for 5 hours targeting different classes HDACs.

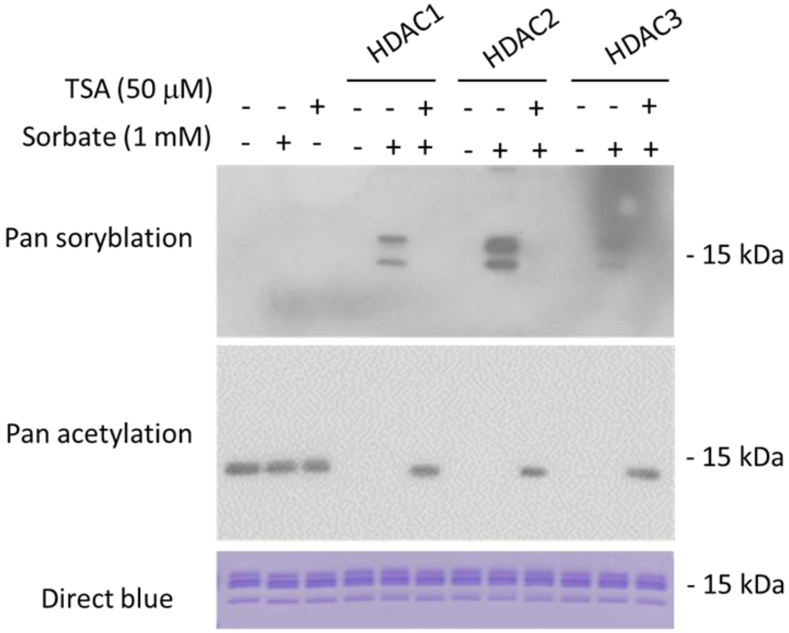


**Fig. S6. In vitro enzymatic assays demonstrating the sorbyltransferase activities of HDAC1-3.** Histones extracted from non-treated 293T cells were incubated with recombinant HDAC1, HDAC2 or HDAC3/NCOR2 with or without sorbate (1 mM) or TSA (50 µM, HDAC inhibitor). After incubation, the protein solution was subjected to western blotting analysis.

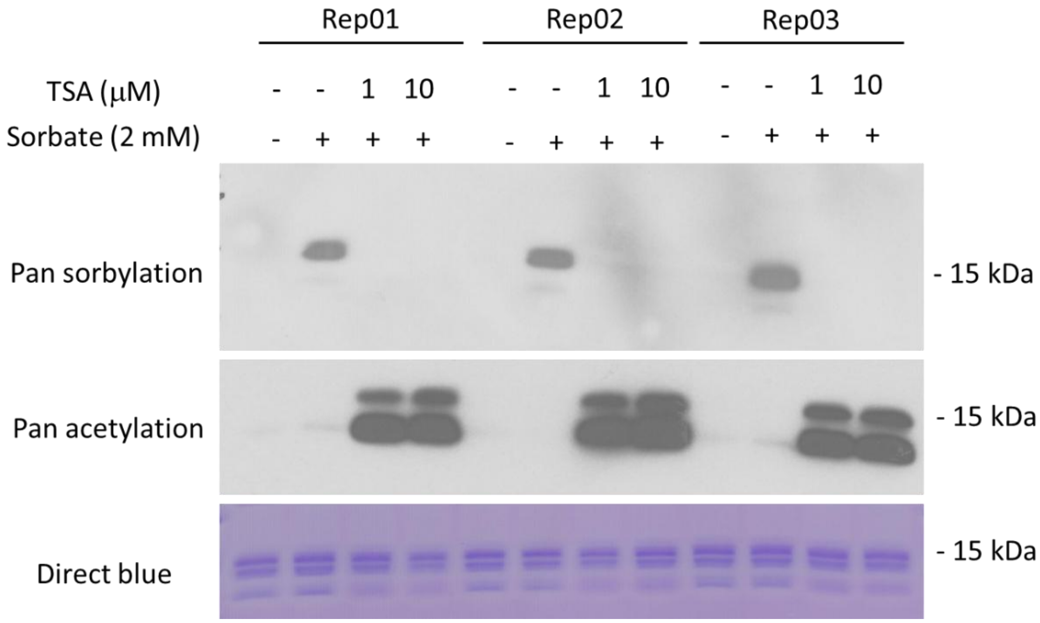


**Fig. S7. Inhibition of sorbylation induction through TSA treatment in 293T cells.** 293T cells were pretreated with TSA (HDAC inhibitor) at indicated concentrations for 30 minutes. Cells with or without TSA pretreatment were supplemented with sorbate for a final concentration of 2 mM and incubated for 6 hours. Finally, the histones were extracted for western blotting analysis with indicated antibodies.

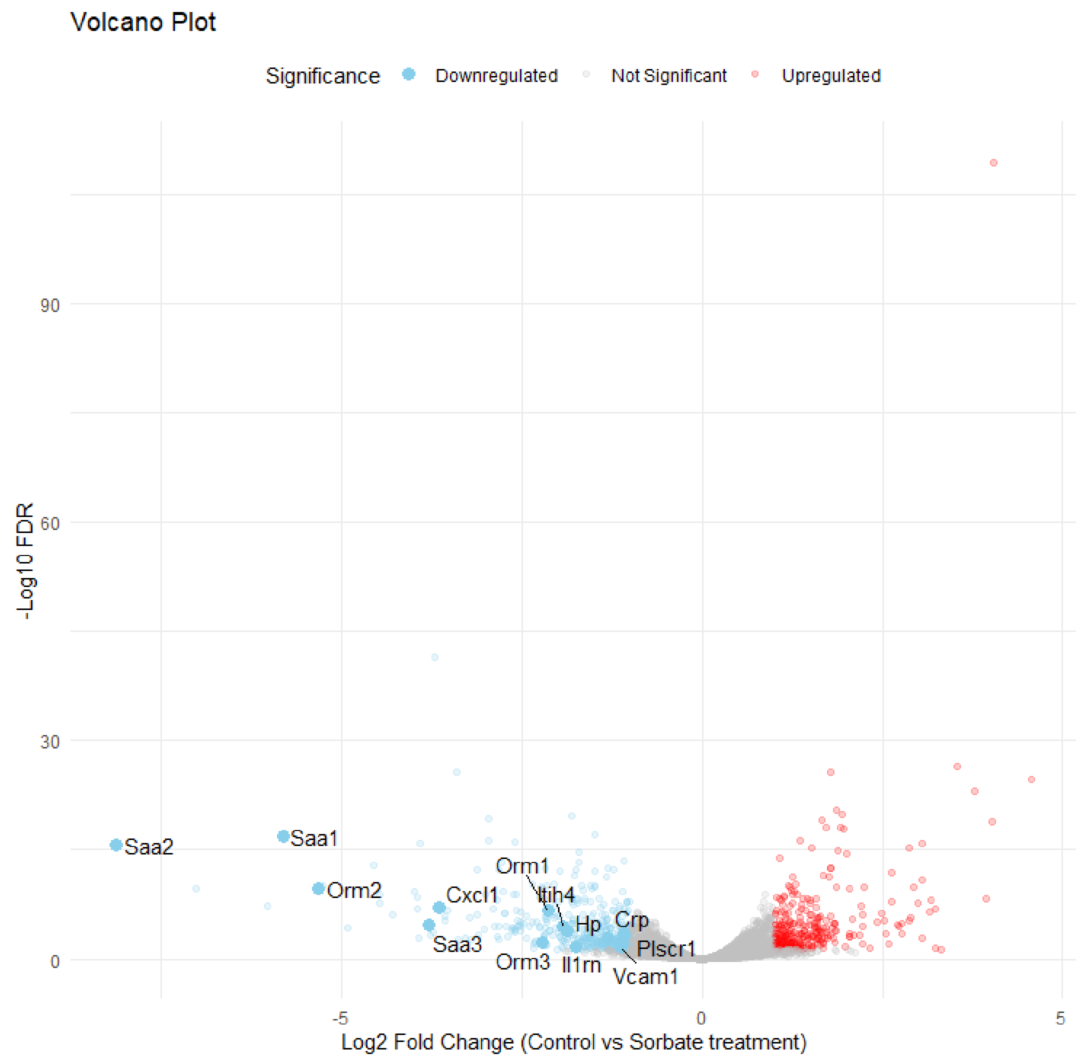


**Fig. S8. The complete volcano plot analysis of gene expression of control and sorbate-treated mouse livers corresponding to Fig. 4A**. Gene expression was analyzed with edgeR (absolute fold change > 2, and FDR < 0.05) and the volcano plot colored genes with significant up-regulation (red dots), downregulation (blue dots) and no significant changes (grey dots).

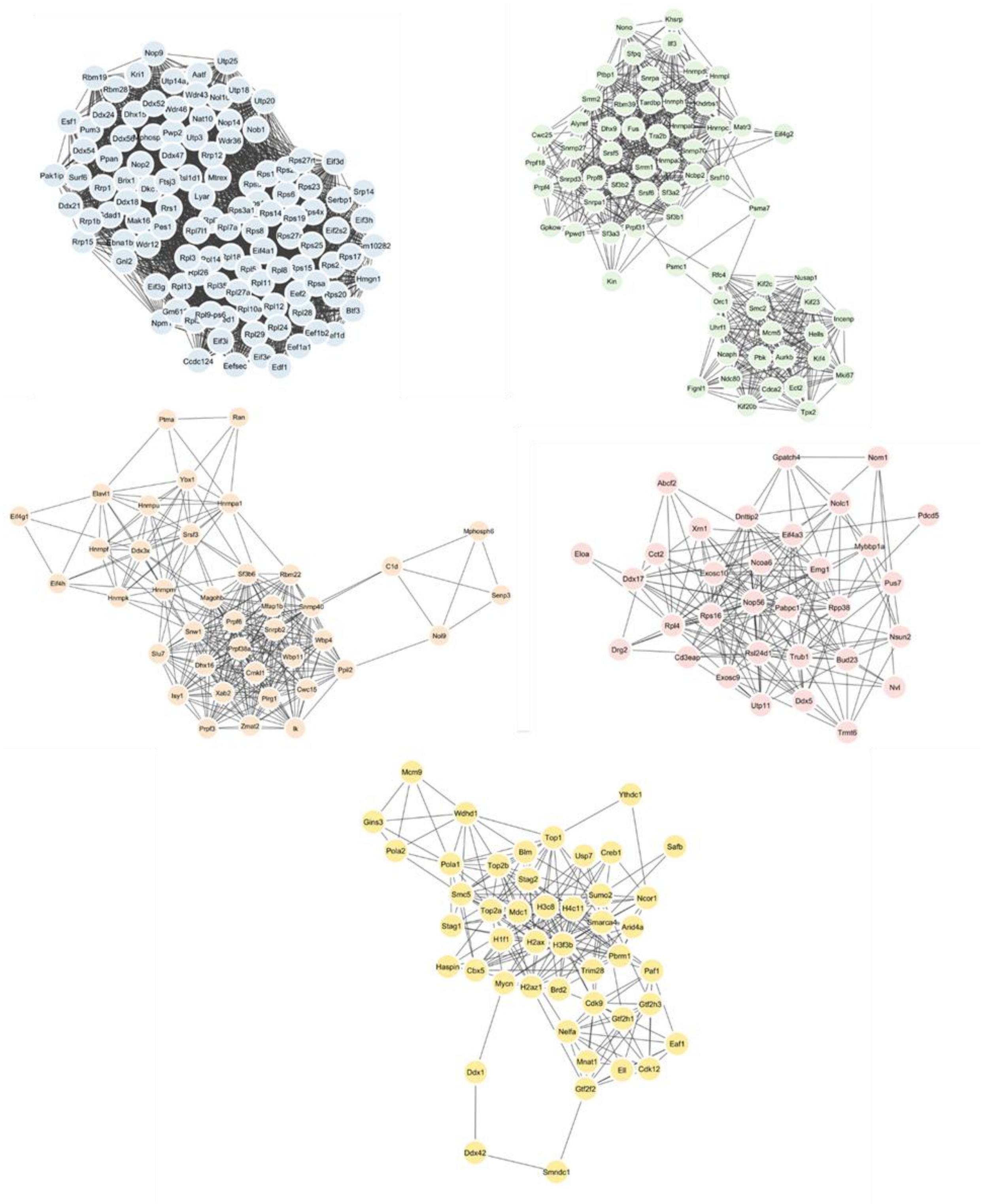


**Fig. S9. Selected subnetworks of Ksor interacting proteins identified from global Ksor protein interaction network by MCODE analysis.**

**Data S1. (separate file)**

A list of identified histone lysine sorbylation sites from in-gel digestion of histones extracted from sorbate-treated (A) 293T cells, (B) RAW264.7 cells and (C) liver of mice fed with diet containing 0.1% or 0.5% potassium sorbate.

**Data S2. (separate file)**

RNA-seq analysis of gene expression dynamics upon sorbate treatment. RNAs were extracted and sequenced from livers of control mice or mice 4 hours after feeding with 0.5% potassium sorbate through oral gavage. (A) A complete list of genes with significant up- or down-regulation from edgeR analysis (FDR < 0.05). (B) A list of significantly down-regulated genes with at least 2 folds decrease. (C) A list of significantly up-regulated genes with at least 2 folds increase. (D) Raw read count from RNA-seq analysis for all samples.

**Data S3. (separate file)**

A list of identified Ksor sites through the immunoprecipitation (IP) and LCMS analysis of RAW264.7 cells with pan anti-Ksor antibody. Raw264.7 cells were treated with 10 mM sorbate overnight and then stimulated with 100 ng/mL LPS for 3 hours prior to lysis, tryptic digestion and IP-LCMS analysis.
